# Joint inference of paired dynamical gene regulatory networks reveals distinct cell-state landscapes of neutrophil reprogramming

**DOI:** 10.64898/2026.09.16.752194

**Authors:** Alex Ren, Yukai You, Mingyang Lu

**Affiliations:** Phillips Exeter Academy, Exeter, NH 03833, USA; Center for Theoretical Biological Physics, Northeastern University, Boston, MA 02115, USA; Department of Bioengineering, Northeastern University, Boston, MA 02115, USA

## Abstract

Disease reprograms cells through changes in gene regulation, yet identifying these changes remains a major challenge. We introduce NetDes-Duo, a computational method that jointly infers transcription factor regulatory network models for two related conditions using scRNA-seq data. The networks are optimized to have minimal topological differences, while the associated ODE models recapitulate single-cell gene expression trajectories for both conditions. On synthetic benchmarks, NetDes-Duo outperformed methods that infer each network independently. NetDes-Duo was applied to neutrophil reprogramming in naive and tumor-bearing mice, and the network-simulated dynamics reproduced the observed cell state transitions. The naive landscape had two well-separated basins, whereas the tumor-bearing landscape was more continuous, with three shallower basins. Perturbation and driving simulations also identified *Cebpb* as a key driver of the tumor-bearing transition, consistent with emergency granulopoiesis literature. We expect NetDes-Duo to be a broadly applicable framework for uncovering the regulatory logic of disease-associated cell state transitions.

## Introduction

A central question in biology is how cells can undergo state transitions in radically different ways, both through the cell’s intrinsic developmental gene regulatory programs and through pathological reprogramming in disease.^1,2^ During development, common progenitors can bifurcate into distinct cell fates, producing cell types with sharply different functions from a shared starting point.^3^ In disease, tumors reprogram surrounding immune and stromal cells toward states that support growth and immune evasion,^4^ while chronic injury redirects normal differentiation toward fibrotic or dysfunctional outcomes.^5^ Neutrophils, for example, were long viewed as short-lived, terminally differentiated cells, yet recent work in myeloid biology has shown that they possess substantial plasticity and heterogeneity, adapting their transcriptional programs to specific tissue microenvironments and pathological states.^6,7^ Under homeostatic conditions neutrophils mature along a trajectory from bone marrow to blood, then traffic to peripheral tissues, such as the lung, to carry out pathogen clearance and tissue surveillance.^8,9^ However, under tumor-bearing conditions systemic and local oncogenic signals hijack this homeostatic trajectory and reprogram neutrophils toward immunosuppressive phenotypes that drive tumor progression, extracellular matrix remodeling, and metastatic seeding.^10,11^ Such reprogramming is fundamentally a change in gene regulation, yet which regulatory interactions are altered and how those changes reshape cellular behavior largely remain unclear.

Traditional experimental workflows for studying cell state transitions during reprogramming often rely on techniques such as complex animal models, conditional knockouts, and extensive ex vivo assays to isolate these transient states, approaches that are labor-intensive, costly, and time-consuming.^12,13^ In recent years, single-cell RNA sequencing (scRNA-seq) has become an increasingly popular technique to study transcriptional programs of cell state transitions.^14^ It captures genome-wide gene expression profiles across cell states along a trajectory, either by profiling cells at multiple time points or by profiling a mixed population of cells at different trajectory stages.^15,16^ Many computational methods have been developed to infer gene regulatory networks (GRNs) from scRNA-seq data using techniques such as statistical inference^17–21^ or deep learning.^22,23^ However, the majority of these methods focus on capturing static networks rather than dynamic changes in GRNs during cell state transitions underlying cell differentiation and disease progression. To address this issue, some recent approaches,^24–28^ including our own previous work^29,30^, have been developed to build non-linear ordinary differential equation (ODE) models of a GRN driving specific cell state transitions, leveraging dynamical systems theory to capture the causal gene regulatory dynamics.

Studying reprogrammed cell-state transitions of the kind described above often requires comparing the regulation of two or more related conditions, such as naive versus tumor-bearing immune cells, untreated versus cytokine-stimulated conditions, or wild-type versus gene-knockout cell lines. However, existing dynamical GRN modeling approaches are largely designed to capture a single trajectory or system, and do not readily support direct comparison of regulatory dynamics across multiple conditions. For example, a straightforward approach is to infer regulatory networks for differing conditions separately. However, regulatory network inference from scRNA-seq data is often unstable, with independent runs on similar datasets retrieving substantially differing GRNs, in terms of both gene nodes and regulatory interaction edges.^31,32^ As a result, we cannot be sure whether differences in two GRNs are biologically grounded or simply artifacts of noise and other errors in the inference process. While some methods such as FSSEM^33^ and sc-compReg^34^ exist to infer a pair of related GRNs, these methods return static networks that describe regulatory differences between conditions, but they cannot model how those differences play out dynamically as cells progress through a state transition. This is an important gap, as the GRNs under two conditions may differ in only a few edges yet produce different gene expression trajectories, different stable states, or different responses to the same signal. Such a model would further enable in silico perturbation simulations, allowing the divergent responses of two conditions to a shared perturbation to be predicted and compared directly.

In this study, we introduce <u>Net</u>work inference and optimization using <u>D</u>ynamical <u>e</u>quation <u>s</u>imulations for <u>Du</u>al-condition <u>o</u>verlap (NetDes-Duo), which jointly infers a pair of transcription factor (TF) regulatory networks together with associated dynamical models capable of characterizing cell-state transitions under two experimental conditions. This work extends our previous NetDes method, which first infers the gene expression trajectories of key TFs, then performs nonlinear ODE fitting and network optimization simultaneously. NetDes-Duo offers two main features. First, by jointly optimizing both networks with a penalty term to minimize the network differences, the resulting pair of GRNs share a common structural topology reflecting conserved gene regulatory programs, while the few condition-specific edges are more likely to represent genuine biological rewiring. Second, the associated ODEs enable in-silico predictions of how cell state transitions differ under these two conditions.

In the following, we first present an overview of NetDes-Duo, with full details provided in the Methods section. We then benchmark the method to show an improvement over the original NetDes method and other popular regulatory network inference methods. Afterwards, we apply the method to neutrophil maturation and differentiation under naive and tumor-bearing conditions to recover key differences in regulatory structure, the landscape of stable states, and response to perturbation between the two conditions. By isolating the TFs and signaling hubs whose regulation differs between conditions, this analysis provides a starting point for designing targeted perturbations intended to reverse pro-tumorigenic neutrophil states.

## Results

### Overview of NetDes-Duo

We introduce NetDes-Duo, a computational method to jointly construct a pair of core TF regulatory networks with high structural similarity across two experimental conditions or bifurcating trajectories. The method is generalized from the previous NetDes method, which allows inference of GRNs and optimization of associated ODE models directly from single-cell gene expression trajectories. The workflow of NetDes-Duo is described in **Fig.1**. First, single-cell gene expression profiles undergo dimensionality reduction, with cells from the two conditions separated out (left half of **Fig.1A**). For each condition, pseudotime is inferred for each cell, and smoothed gene expression time trajectories are obtained by LOESS regression. These time trajectories are then grouped into gene clusters based on their overall shapes (middle panel of **Fig.1A**). From each cluster, over-representation enrichment analysis is performed against literature-curated TF target gene sets, from which core TFs are identified. From the analysis, we identify core TFs shared between conditions, along with a few condition-specific TFs (rightmost panels in **Fig.1A**). These core TFs are then connected to form a pair of initial TF regulatory networks using literature-based and motif-derived TF-target databases (TRRUST^35^ and RcisTarget^18^). Starting from the initial networks, we follow the original NetDes procedure for network optimization and nonlinear ODE fitting (see the Methods), with one key modification as follows. NetDes-Duo assumes that both conditions correspond to similar underlying GRNs. Therefore, the pair of GRNs is optimized with minimum edge differences by introducing a penalty term, as described in detail in the Methods (**Fig.1B**). Finally, using the optimized dynamical models and GRNs, we can perform analyses such as identifying common and condition-specific regulatory interactions, comparing regulatory landscapes, and running gene expression dynamics simulations under driving signals or gene perturbations (**Fig.1C**).

**Fig 1.**
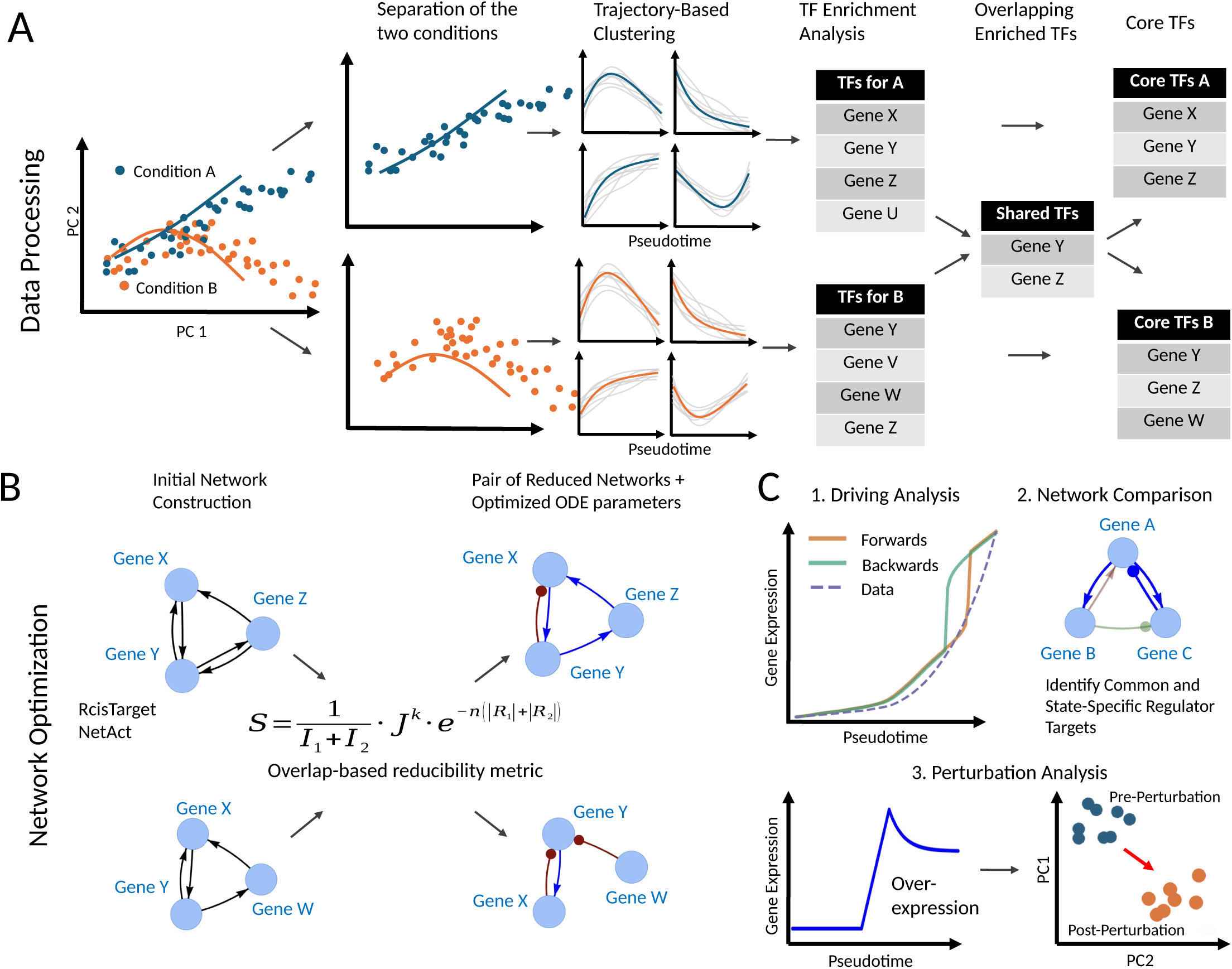
Overview of workflow for NetDes-Duo. **(A)** scRNA-seq data from cell state transitions are first separated by condition. For each condition, pseudotime is inferred, and smoothed time trajectories are obtained for each gene. Gene clustering is then performed on these gene trajectories based on overall shape, followed by over-representation analysis to infer core TFs. The final core TF list contains the top ranked TFs, both shared and condition-specific. **(B)** Two initial TF regulatory networks are first built by connecting two TFs according to literature-based and binding-motif-based transcriptional regulatory interactions. Using information from both initial networks, NetDes-Duo constructs two highly overlapping optimized networks along with the associated ODE models. **(C)** The optimized network models can be used to perform (1) forward and backward driving simulations, (2) identification of common and path-specific transcriptional regulatory interactions, (3) stochastic simulations to characterize the cell state landscape, and (4) perturbation simulations to identify key TFs that drive state transitions.

### Benchmarking with synthetic regulatory networks

To benchmark NetDes-Duo against existing regulatory network inference methods, we constructed 20 test cases, each containing a pair of synthetic regulatory networks with high structural similarity (each pair with Jaccard index >0.7; an example is shown in **Fig.2B**, with complete network trajectories shown in **Fig.S1**). Each constructed network contains the same seven genes (TFs) and 18 interactions, half activating and half inhibiting. Each network consisted of one driving signal (e.g., blue trajectory in **Fig.2A**), while the rest of the trajectories were generated via ODEs with randomly sampled kinetic parameters (more details in Methods). Due to the differences in both network topology and kinetic parameters, gene expression trajectories in the paired networks are also different (**Fig.2A**).

**Fig 2.**
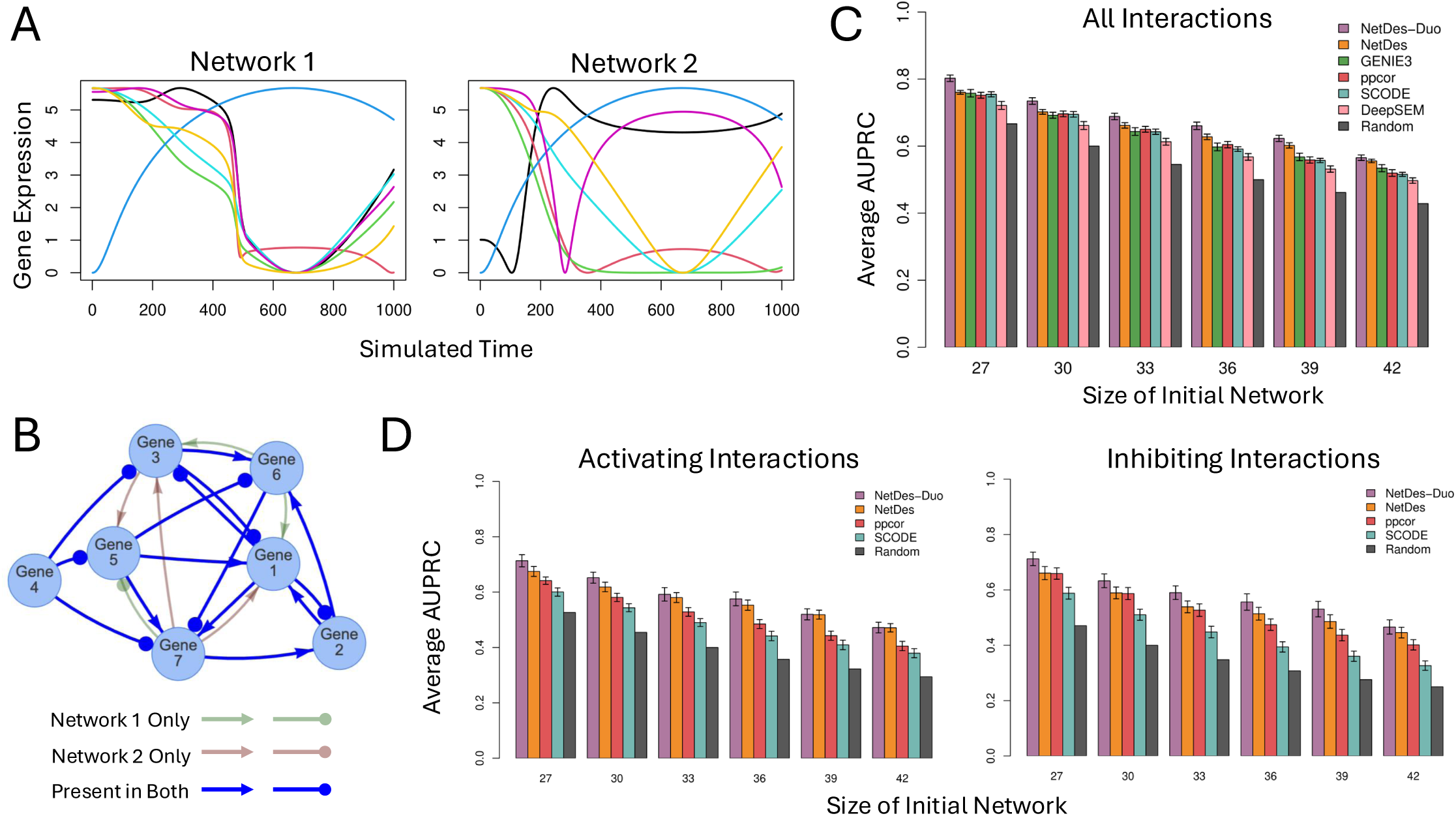
Benchmarking using synthetic gene expression trajectories. **(A)** Example of gene expression trajectories for a pair of synthetic networks. The blue curve represents the gene expression trajectory of the driving signal, shared across both networks. The other colors represent the gene expression trajectories of the other network genes, simulated by Hill-type ODEs according to the network topologies with randomly sampled parameters. **(B)** Diagram of the topologies of the two networks for the example in panel A. Lines with arrowheads indicate activation, while lines ending in dots indicate inhibition. Edge colors indicate whether an edge is presented in network 1 only (green), network 2 only (red), or both (blue). **(C)** Mean AUPRC for each network construction method on a benchmark set of 20 network pairs. Each initial network contained 18 ground-truth edges and 9-24 decoy edges, resulting in a total of 27 to 42 edges per network. **(D)** Mean AUPRC computed separately by activating (left panel) and inhibiting (right panel) interactions.

Next, we constructed initial networks by adding 9-24 decoy edges on top of the 18 ground-truth interactions, resulting in network sizes between 27 and 42. NetDes-Duo was then applied on each pair of networks, while all other methods were applied on each of the networks separately. Specifically, we varied the NetDes-Duo parameters (more details in Methods) to optimize GRNs of varying sizes, which we used to calculate the Area Under the Precision-Recall Curve (AUPRC) per network. We found that NetDes-Duo outperformed all other methods (**Fig.2C**) in mean AUPRC across all initial network sizes. In addition, consistent with the design of the paired-network objective, NetDes-Duo recovered optimized network pairs with substantially greater overlap than any other applied method (**Fig.S2**). When considering the regulatory sign (i.e., the activating/inhibiting nature), NetDes-Duo also obtained the highest AUPRC scores when considering activating and inhibiting interactions separately (**Fig.2D**). The base NetDes method also performed strongly, ranking second or third across all initial network sizes and interaction categories, including all interactions, activation-only, and inhibition-only (**Fig.2C-D**). The high performance of both NetDes and NetDes-Duo benefits from the incorporation of dynamical models into network inference. Importantly, NetDes-Duo consistently outperformed the base NetDes across all tests, demonstrating that explicitly incorporating similarity between paired networks provides a measurable performance advantage in the inference of these structurally similar networks.

### Application to Neutrophil Reprogramming in Naive and Tumor-Bearing Mice

Next, we applied NetDes-Duo to a scRNA-seq dataset on neutrophil reprogramming. Specifically, we examined scRNA-seq data for neutrophils collected from naive mice and from tumor-bearing mice across three organ types (bone marrow, blood, and lung).^36^ These three organ types represent state transitions across neutrophils’ entire life cycle, which generally progresses from bone marrow (BM) to blood and then to lung. When the processed scRNA-seq data for both conditions were projected onto the first two principal components (PCs) (**Fig.S3**), a clear bifurcation of state transitions arose, with a common BM neutrophil state diverging into distinct lung-associated cell states. However, since these two different neutrophil state transitions come from fundamentally the same underlying biology under different environmental conditions, we expected that the underlying regulatory networks controlling maturation and differentiation would be highly similar. Thus, we applied NetDes-Duo by first constructing two separate initial TF regulatory networks using the condition-specific gene expression data and then optimizing a pair of GRNs and associated ODEs.

To begin, we applied principal component analysis (PCA) to the expression of the top 500 highly variable genes (HVGs) (projection onto the first two PCs shown in **Fig.3A**). We observed continuous progression of neutrophil maturation and tissue-specific reprogramming from bone marrow to blood to lung, common to both naive and tumor-bearing conditions. For each condition, we inferred pseudotime using a diffusion map via the Destiny package,^37^ which was used to infer gene expression trajectories by LOESS regression for approximately 3000 highly variable and highly expressed genes. These genes were first automatically clustered into 15 naive and 14 tumor-bearing gene clusters based on overall shape (**Figs.S4, S5** and the Methods). These initial clusters were then manually combined into 6 large gene clusters for the naive condition and 7 large clusters for the tumor-bearing condition (**Fig.3B**). Then, we performed enrichment analysis (using Fisher’s exact test) to identify TFs whose targets were enriched in one of the clusters.

**Fig 3.**
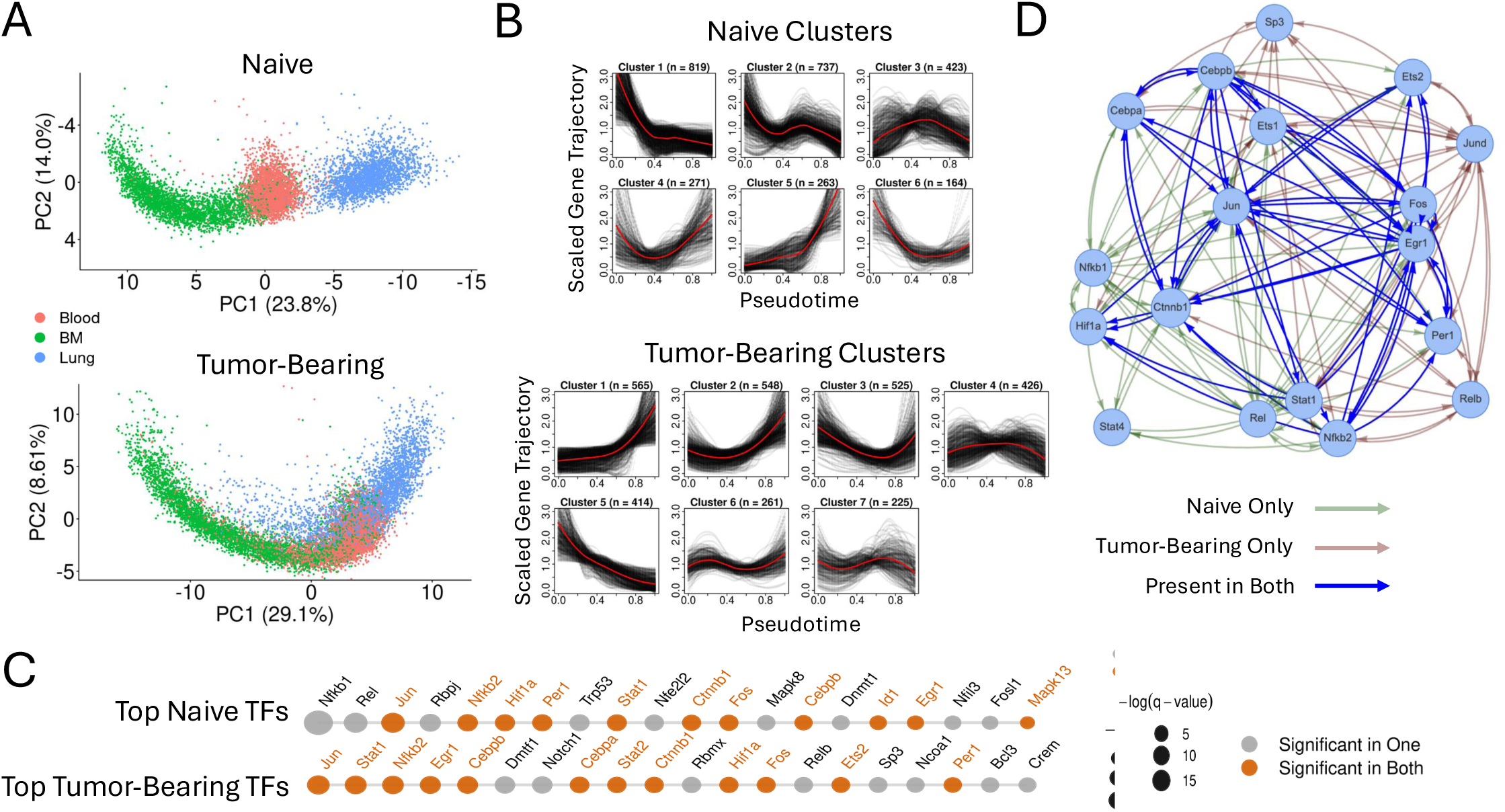
Cell state transitions observed in neutrophil reprogramming from naive and tumor-bearing mice. **(A)** Projection of single-cell expression of neutrophils onto their first two principal components. Data for the naive and tumor-bearing mice are shown in the top and bottom panels, respectively. Cells are represented as points colored by tissue source (blood, bone marrow, and lung). **(B)** 3000 genes that were both highly variable and highly expressed are clustered based on the overall shape of their trajectories. Six clusters were identified for the naive case (top panel), while seven were identified for the tumor-bearing case (bottom panel). Each gene’s expression curve is scaled to have an average of one, and the red line represents the average expression level of the cluster. **(C)** Top 20 TFs were identified for the naive (first row) and tumor-bearing (second row) cases by applying over-representation analysis to each gene cluster in panel B. TFs are ranked from left to right by q-value. TFs with q-value < 0.25 in both naive and tumor-bearing lists are colored in orange. **(D)** Diagrams of both initial TF regulatory networks derived from the naive and tumor-bearing core TF lists. Blue lines indicate an interaction found in both networks. Green and red lines indicate an interaction only found in the naive and tumor-bearing networks, respectively. At this stage, all interactions are unsigned with respect to activation/inhibition.

From the enrichment analysis (**Fig.3C**), we identified 15 shared TFs across the naive and tumor-bearing conditions. After utilizing RcisTarget and TRRUST to build an initial network, we observed that only 12 (*Cebpa*, *Cebpb*, *Ctnnb1*, *Egr1*, *Ets1*, *Ets2*, *Fos*, *Hif1a*, *Jun*, *Nfkb2*, *Per1*, *Stat1*) of these 15 TFs were well connected in both cases. We then incorporated three top state-specific TFs to each case using a selection metric that weighted both q-value from the enrichment analysis and high connectivity with the rest of the network (more details in the Methods). Specifically, *Rel*, *Nfkb1*, and *Stat4* were chosen for the naive condition and *Relb*, *Jund*, and *Sp3* were chosen for the tumor-bearing condition. Finally, applying RcisTarget and TRRUST again to these refined core TFs resulted in a pair of initial networks with 15 TFs for each condition (**Fig.3D**).

These TFs are well-established regulators of neutrophil biology. *Cebpa* is the principal regulator of steady-state granulopoiesis^38,39^ whereas *Cebpb* sustains neutrophil production during emergency granulopoiesis, which is activated under stress.^40,41^ Beyond the C/EBP family, the AP-1 components *Fos* and *Jun* participate in the inflammatory transcriptional programs of myeloid cells and of tumor-associated neutrophils.^42,43^ *Hif1a* acts later in the neutrophil lifecycle, sustaining survival at hypoxic inflammatory sites through NF-κB-dependent activity.^44^ Similarly, *Per1* belongs to the circadian timer that paces neutrophil aging and egress.^45,46^ Meanwhile, Ctnnb1 is transcriptionally upregulated in neutrophils after ischemic injury, where it promotes survival.^47^ Finally, Egr1, Nfkb2, and Stat1 relate more broadly to stress- and inflammation-responsive programs.^48–50^

The condition-specific TFs were also biologically interpretable. In the naive network, *Rel* and *Nfkb1* play roles in canonical NF-κB-associated inflammatory regulation, whereas the tumor-bearing network contains *Relb*, a core component of noncanonical NF-κB signaling.^43,51^ This separation is consistent with prior evidence linking *Relb*-positive neutrophil programs to tumor-associated inflammatory states, in which *Relb* was identified as the central transcriptional regulator of a pro-metastatic neutrophil subset.^52^ Thus, the TF selection procedure recovered both a shared maturation-associated regulatory program and condition-specific regulators with links to tumor-associated neutrophil reprogramming.

### Constructing Condition-specific Neutrophil GRNs

NetDes-Duo was next applied to the pair of initial networks, aiming to obtain highly overlapping sets of regulators for every shared network gene. NetDes-Duo has two hyperparameters related to the penalty term: deletion parameter n and overlap penalty parameter k. Hyperparameter tuning on the current application showed that fitting error, in terms of the average mean squared error (MSE), increased sharply when the deletion parameter exceeded n = 0.5, corresponding to inferred networks of approximately 40-50 edges (**Fig.S6A, B**). Varying the overlap penalty parameter showed that k = 1.5 maintained relatively low MSE while preserving network similarity (**Fig.S6C**). These hyperparameter values were therefore used for subsequent analyses.

After the GRN optimization, the pair of TF regulatory networks were substantially smaller than the initial networks while retaining strong structural similarity between conditions. The optimized networks contained 41 interactions in the naive condition and 43 interactions in the tumor-bearing condition, compared with 101 and 99 interactions in the corresponding initial networks (the two optimized networks are shown individually in **Fig.S7**). As expected, these optimized networks showed significantly higher overlap than pairs of inferred networks from other methods (**Fig.S9**). The optimized ODE models fit the time trajectories well across all TFs (**Fig.4C**; full fitting results in **Fig.S8**), indicating that the reduced networks retained sufficient regulatory information to reproduce the observed gene expression dynamics. Of note, the networks included the TFs whose time trajectories were very similar across the two conditions (such as *Jun*) and TFs whose trajectories differed drastically (such as *Stat1*).

**Fig 4.**
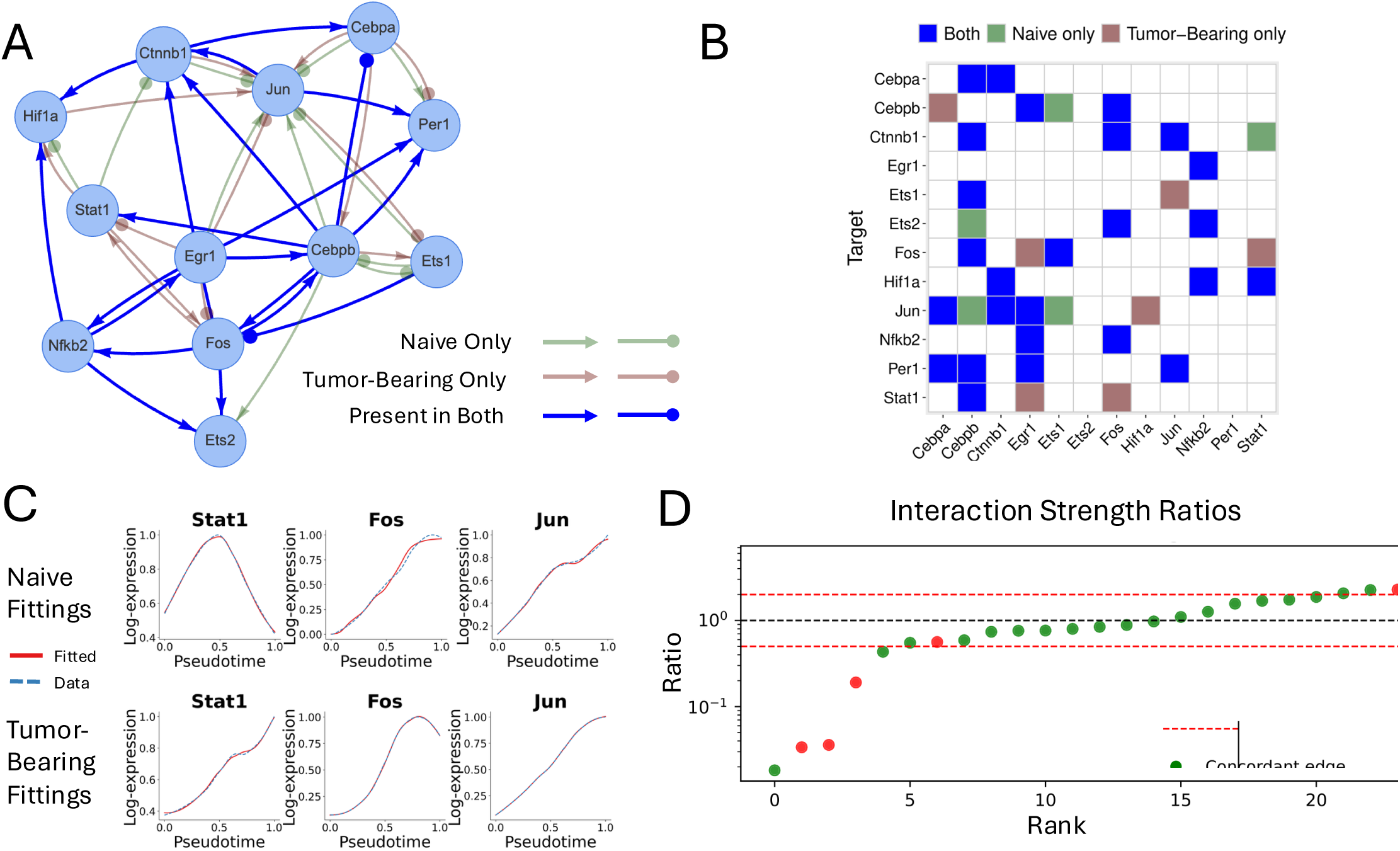
Optimization of paired networks for naive and tumor-bearing conditions. **(A)** Diagrams of the optimized naive and tumor-bearing networks. Lines with arrowheads represent activation, and lines ending in dots represent inhibition. Edge colors indicate whether an edge is presented in naive network only (green), tumor-bearing network only (red), or both (blue). **(B)** Comparison of TF-TF interaction matrices between the naive and tumor-bearing networks. Blue squares represent an interaction present in both networks, green squares for naive network only, red squares for tumor-bearing network only. The matrix does not indicate sign (activation vs. inhibition) and is constrained to the 12 shared TFs between the networks. **(C)** Gene expression trajectories for the scRNA-seq data (blue dotted line) and the optimized ODE model (red solid line) for selected genes. Complete trajectory fitting results for all TFs are shown in **Fig.S8**. **(D)** Ratio of interaction strengths (interaction strength defined by the maximum fold changes for each interaction) between the overlapping edges in the naive and tumor-bearing networks. Green dots represent edges with consistent activation/inhibition between the two networks, while red dots represent sign disagreements. The ratios are plotted on a log scale, and the dotted red lines represent ratios of 0.5 and 2.

Among the twelve overlapping TFs, we identified 26 shared TF-TF interactions (**Fig.4A, B**). Of these interactions, 20 had concordant regulatory signs between conditions, while six differed in sign. In addition, the optimized networks retained five naive-specific and seven tumor-bearing-specific interactions amongst these twelve TFs (**Fig.4A, B**). Among these differences, *Jun* had the most changes in its regulators: *Jun* had two naive-only regulations, one tumor-bearing-only regulation, and all three shared regulators switched sign between the two conditions. The largest regulatory shift for *Jun* between naive and tumor-bearing neutrophils could be due to its role as the AP-1 hub that integrates MAPK signaling^42^ from multiple tumor-associated stimuli, such as cytokines, ROS, and hypoxia.^53,54^

In addition, we identified *Cebpa*, *Stat1*, and *Cebpb* as the regulators which played significantly different roles in the two conditions. In particular, *Cebpa*’s and *Stat1*’s outgoing edges were entirely disjoint between conditions after considering signs, and a large share of *Cebpb*’s interactions also changed. *Cebpa* gained a tumor-bearing-only target and switched the sign of both its shared targets, while *Stat1* showed one sign swap, one naive-only edge, and one tumor-bearing-only edge. Several of the differentially wired regulators have established roles in both steady-state and emergency granulopoiesis. *Cebpa* is the key regulator of steady-state granulopoiesis, and its loss blocks neutrophil differentiation and G-CSF receptor signaling. ^38,39^ In contrast, emergency granulopoiesis, which is activated under infection and other stress conditions, depends instead on *Cebpb*, the key factor induced by G-CSF-STAT3 signaling that sustains granulocyte production in *Cebpa*-deficient progenitors.^40,41^ Beyond granulopoiesis, *Cebpb* is also required for the immunosuppressive activity of tumor-induced myeloid cells.^55^

One other interesting change we noticed involved *Jun* and *Ctnnb1*. These TFs formed a two-node loop in both conditions, consistent with reported reciprocal regulation between β-catenin and JNK/c-*Jun* signaling.^56^ *Jun* activated *Ctnnb1* in both networks, but *Ctnnb1*’s regulation of *Jun* reversed sign, so that the loop constituted negative feedback in the naive condition and positive feedback under tumor-bearing conditions. Since changes in feedback sign can alter the qualitative dynamics of a circuit, this localized rewiring illustrates how a small number of edge-level changes may reshape network behavior between related conditions.

Finally, we compared the fitted strengths of the shared interactions between two GRNs. For each of the 26 shared edges, we computed the ratio of its interaction strength (defined as the maximum fold-change parameter in the ODE model) in the naive network to that in the tumor-bearing network (**Fig.4D**). Here, an interaction strength much less than one represents strong inhibition and a value much greater than one represents strong activation. Most edges, and concordant edges in particular, had ratios within roughly a factor of two, indicating that shared edges also tended to retain a similar quantitative effect on their targets. As expected, the sign-discordant edges were concentrated at the extremes of the ratio distribution, so that the interactions that reversed sign were also among those whose fitted strengths diverged most. The conserved strengths of the shared edges reinforce the view that the two conditions are governed by a largely common regulatory backbone, with differences manifesting in a small set of condition-specific and/or sign-switched interactions.

### Stochastic Simulation Reveals Divergent Naive and Tumor-Bearing Landscapes

After constructing the condition-specific GRNs, we analyzed the dynamical behavior of the GRNs in establishing the gene expression states and the regulation of the transitions between states using a dynamical systems approach. Here, we performed stochastic simulations by adding Gaussian noise terms to the optimized ODEs, starting from 5000 random initial conditions of gene expression. Each simulation could be regarded as representing a simulated cell. Simulations were run for sufficiently long so that the system reached a nearby steady state. PCA was then performed on the final gene expression profiles for all simulated cells, and a principal curve was fitted through the resulting distribution of points. Each cell was then assigned a principal curve coordinate, its position along this curve, running from 0 at the immature end to 1 at the mature end (**Fig.5A**; see Methods for more details).

**Fig 5.**
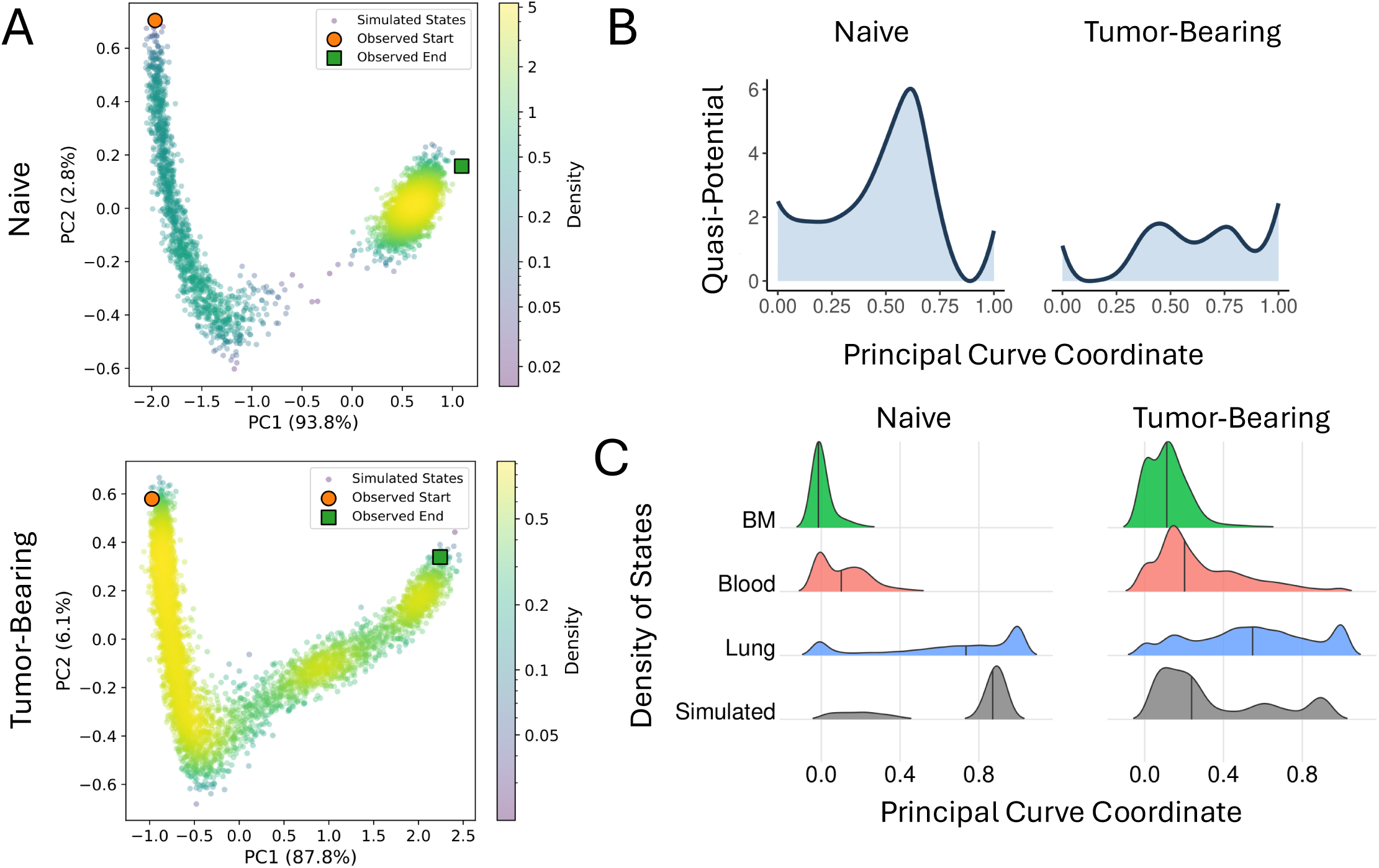
Stochastic simulations capture condition-specific neutrophil cell state landscapes. **(A)** Projection of simulated gene expression profiles (blue dots) onto the first two principal components. Stochastic simulations were performed using the optimized ODE models with added noise terms for both the naive (top panel) and tumor-bearing (bottom panel) conditions. Orange and green circles represent the starting and ending gene expression profiles from the scRNA-seq data, respectively. **(B)** Quasi-potential along the principal curve, derived from the PC1 vs. PC2 plots in panel **A**. **(C)** Histogram of cells in each tissue (bone marrow in green, blood in red, and lung in blue) and from the simulation (grey) along the principal curve (derived from the simulation). Left and right panels show the results for the naive and tumor-bearing conditions.

Interestingly, both stochastic simulations recovered gene expression distributions spanning the full range of the experimental gene expression states along the principal curve, as shown in **Fig.5A** and **Fig.3A**. In both the PCA of the simulated data and the PCA of the scRNA-seq data, the naive case formed two relatively distinct clusters, one containing neutrophils from the bone marrow and blood and the other containing neutrophils from the lung. Meanwhile, the tumor-bearing case formed a much more continuous trajectory with 3 smaller and less distinctly separated clusters. Notably, the ODE models were optimized against smoothed expression trajectories without learning the stability and density of steady states, but the models still captured the number and separation of cell states observed in the data.

Next, we calculated the quasi-potential of the landscape along the principal curve as *U* −log (*P*), such that high-density regions correspond to potential wells (**Fig.5B)**. The naive landscape resolved into two wells separated by a pronounced barrier, one at the immature and one at the mature end. Notably, the immature well was shallower than the mature well, so the landscape favors forward progression from the immature to the mature state. Meanwhile, the tumor-bearing landscape resolved into a more continuous trajectory. Specifically, we observed three wells separated by significantly lower barriers, with every tumor-bearing inter-well barrier being less than half the height of the naive barrier. This suggests that under the tumor-bearing condition, state transitions during neutrophil reprogramming would be more easily induced by signaling.

To examine how the measured cells from each tissue relate to these simulated landscapes, we projected the original single-cell data onto the principal curve derived from the corresponding simulation for each condition **(Fig.5C, Fig.S10A**). To further analyze this, we binned cells along the simulated principal curve and examined the tissue composition of each bin (**Fig.S10B**). In both conditions the earliest bins were dominated by bone marrow neutrophils, the middle bins by blood neutrophils, and the latest bins almost entirely by lung neutrophils, so that the three tissues were laid out in sequence along the inferred trajectory. This progression matches the expected maturation order and is consistent with our idea of the cell-state transition between the organs.

**Fig.5C** also shows the histogram of the neutrophils along the principal curve, separated by organ type. In the naive condition, bone marrow and blood neutrophils landed almost entirely in the first, early-progression cluster. The neutrophils from blood formed two adjacent humps, but both fell within this single cluster. Lung neutrophils were distributed across the trajectory, with a low, broad shoulder at earlier progression and a peak in the second, late-progression cluster. In the tumor-bearing condition, bone marrow and blood neutrophils once again peaked early, though a substantial fraction of blood cells extended well beyond the first cluster. Lung neutrophils formed a far more continuous distribution than their naive counterparts, spread broadly across the trajectory. However, we still observed three peaks corresponding to the three simulated peaks at early, intermediate, and late progression. The correspondence between where the measured cells fall and where the simulated landscape places its basins suggests that the recovered wells reflect features of the observed cell-state distribution, further validating the optimized network and the associated dynamical models.

### Perturbation and Driving Simulations Identify Condition-Specific Drivers of Neutrophil Reprogramming

With the established dynamical models, we examined the role of different genes and their combinations in driving the cell state transitions for each condition. Starting from states sampled from each condition’s baseline steady-state cloud (i.e., untreated condition), we applied a single impulse overexpressing an individual TF or a pair of TFs, continued the stochastic simulation for a fixed relaxation window of 2,000 steps, and scored each perturbation by its change in mean progression along the principal curve (**Fig.6A**; see Methods). Larger mean values indicate more progression toward the mature state, while smaller mean values indicate less progression from the immature (BM) state. Note the reported shifts from these simulations are finite-time displacements; because the impulse is applied at a single time point and the underlying models are unchanged, these displacements do not represent a new equilibrium distribution.

**Fig 6.**
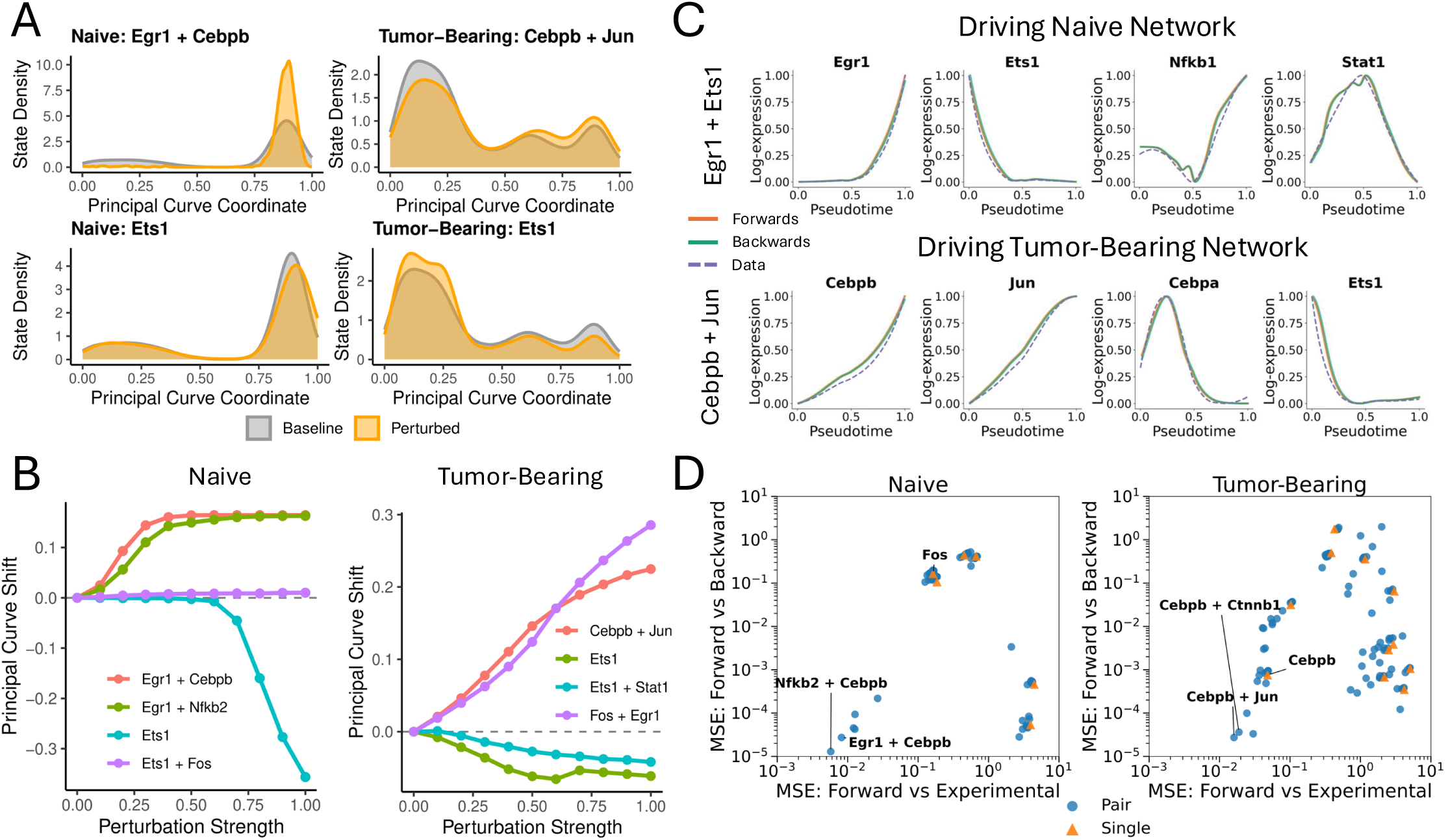
Driving and perturbation simulations identify condition-specific driver genes. **(A)** Transient over-expression was simulated for a single TF or pair of TFs starting from the simulated gene expression profiles from the untreated condition (as shown in Fig.5A). Each panel shows the histogram of cells along the principal curve at the end of the relaxation window, for over-expression of *Egr1* and *Cebpb* for the naive condition (top left), over-expression of *Cebpb* and *Jun* for the tumor-bearing condition (top right), over-expression of *Ets1* for the naive condition (bottom left), and over-expression of *Ets1* for the tumor-bearing condition (bottom right). Grey and orange histograms show the baseline (untreated condition) and perturbed conditions, respectively. **(B)** Average shift in progression along the principal curve under the overexpression perturbations is shown with respect to the strength of the transient over-expression for selected TF perturbations in both conditions. **(C)** TF pairs driving the network gene expression changes were identified in both conditions. Results are shown for the top pairs in each condition: *Egr1* + *Ets1* (naive), and *Cebpb* + *Jun* (tumor-bearing). The plots show the gene expression trajectories for selected TFs under forward driving (solid red lines), backward driving (solid green lines), and the observed scRNA-seq data (dotted blue lines). Trajectories for all TFs in the network are shown in **Fig.S11**and **S12**. **(D)** Scatter plot of driving quality (MSE between the forward driven curve and the experimental data) against hysteresis (MSE between the forward and backward driven curves). Blue dots represent driving with pairs of TFs, while orange triangles represent driving with a single TF. The tumor-bearing condition shows a much larger number of TFs and TF pairs that can drive the system well.

The top forward pair (producing the greatest increase in mean progression) was *Egr1* and *Cebpb* for the naive case and *Cebpb* and *Jun* for the tumor-bearing case (the full ranking of single-TF and paired perturbations for the naive and cancer cases are given in **Table S1** and **Table S2**). In both conditions, essentially every top-ranked pair contained a single dominant TF; this was *Egr1* in the naive case and *Cebpb* in the tumor-bearing case. Over this window, these dominant TFs produced the largest movement toward the mature state in each condition. Notably, *Cebpb* was identified as one of the top genes in both conditions, serving as the leading partner to the dominant driver *Egr1* in the naive case and as the dominant driver itself in the tumor-bearing case. When increasing the perturbation strength of each of these top pairs (**Fig.6B**), we observed a noticeable shift in mean progression in both the naive and tumor-bearing cases even at low perturbation strengths. The shift in the naive condition does level off past a certain strength level, although this is because the perturbation had by then moved essentially all the simulated cells into the second, mature well.

Backward movement along the trajectory was driven primarily by *Ets1* in both conditions, with *Ets1* overexpression alone outperforming any pair of TFs (**Fig.6A**). However, the two networks differed sharply in how readily this backward movement could be induced (**Fig.6B**). In the naive network, we observed a threshold-like curve, where essentially no movement happened until the perturbation strength was raised to a sufficiently high level, after which cells began to move strongly backward. This is consistent with our earlier results modeling a bistable switch, where the mature state is stable and difficult to transit out of. In the tumor-bearing network, by contrast, *Ets1* drove cells backward even at low perturbation strength, reflecting the more continuous landscape with shallower wells. Interestingly, however, the tumor-bearing response in mean progression was not strictly decreasing as the perturbation strength for *Ets1* increased (**Fig.6B**). The different responses to the perturbation are mainly due to the differences in the underlying regulatory landscapes of neutrophil reprogramming, and partly because the mature state is dominant without perturbation in the naive condition, while the immature state is dominant without perturbation in the tumor-bearing condition.

Next, we performed driving simulations by clamping a single TF or a pair of TFs to their inferred gene expression trajectories and used this as a signal to drive the rest of the network according to the derived ODE models. For each single TF or pair of TFs, we recorded how well the driven trajectories matched the data (forward-versus-actual MSE) and how much the forward and backward drives disagreed (forward-versus-backward MSE), the latter measuring hysteresis. Low hysteresis implies a lower barrier between states, while pronounced hysteresis suggests a more rugged landscape that makes transitions harder to achieve.^29^

In the naive condition, driving the single top TF, *Egr1*, failed to cleanly reproduce the transition: forward and backward driving diverged, and several TFs showed poor agreement with their observed trajectories in scRNA-seq, indicating that a single driver could not carry the system across the transition (**Fig.S11A**). Driving *Egr1* together with *Ets1* resolved this failure, as forward and backward trajectories converged and agreed well with the trajectories from the data across all other TFs (selected trajectories in **Fig.6C**, complete trajectories in **Fig.S11B**), indicating that a second driver was required to traverse the state transition in the naive condition. Consistent with this, low-error, low-hysteresis driving solutions were scarce in the naive network and all required multi-TF combinations (**Fig.6D**).

In comparison, cell state transitions were much easier to drive in the tumor-bearing condition. A single TF, *Cebpb*, was sufficient to drive the transition and already reproduced the trajectories at a qualitative level without pronounced hysteresis (**Fig.S12A**). Adding a second TF, *Jun*, only refined the fit without changing this qualitative behavior (**Fig.6C**, complete trajectories in **Fig.S12B**). More broadly, the tumor-bearing network yielded far more successful driving solutions than the naive network; we observed a large population of single- and pair-driving choices achieving low forward-versus-backward disagreement (**Fig.6D**). Together, these results suggest that the tumor-bearing microenvironment may reshape the regulatory landscape to be more permissive to state transitions.

## Discussion

In this work we introduced NetDes-Duo, a computational method that infers two condition-specific TF regulatory networks along with a fitted set of nonlinear ODEs using single-cell gene expression data. The motivation for inferring a pair of regulatory networks is to bring additional biological constraints to bear on an inference problem, assuming that the regulatory program is largely conserved across conditions. Specifically, network inference from scRNA-seq is unstable as independent runs on similar datasets can return substantially different networks. By incorporating the relevant biological knowledge that two conditions describing the same system should have similar regulatory networks, NetDes-Duo shows improved performance and stability in network inference. NetDes-Duo constrains only network structure and allows the two conditions to differ in their fitted ODE parameters, so it can be applied even when the same set of TFs is governed by different dynamics. We evaluated the method on 20 pairs of synthetic networks with varying numbers of added decoy edges and then applied it to neutrophil reprogramming from naive and tumor-bearing mice. On the synthetic benchmarks, NetDes-Duo achieved the highest AUPRC of all methods tested at every proportion of decoys. The increased performance is likely due to the two networks providing doubled opportunities for the optimization algorithm to escape local optima. Since network optimization for one condition informs the other, only one of the two searches needs to find a good solution to improve the optimization for the whole pair. Finally, NetDes-Duo was applied to the neutrophil data to construct the TF regulatory networks for both the naive- and tumor-bearing conditions. Between the two networks, we observe a strong common backbone as most of the shared interactions have similar regulatory effects (within roughly a factor of two) in the fitted ODEs for the two conditions. In addition, the simulated PCA clouds obtained from the fitted ODEs recapitulated the cluster structure of the original scRNA-seq data in both conditions, and projecting the measured neutrophil data onto each simulated principal curve ordered them from bone marrow to blood to lung along the curve as expected.

The dynamical systems modeling elucidates the similarity and differences in the gene regulation of neutrophil reprogramming in both the naive and tumor-bearing conditions. In terms of the landscape of the cell states along the reprogramming trajectories, according to the ODE models, the naive landscape resolved into two wells separated by a pronounced barrier, whereas the tumor-bearing landscape resolved into three wells whose inter-well barriers were each less than half the height of the naive barrier. Additionally, the naive landscape contained a significantly shallower immature well than its mature well, meaning maturation is close to one-way under homeostasis.^57,58^ This collapse of the potential barriers in the tumor-bearing condition is biologically reflected in the rapid egress of pathologically rewired, immunosuppressive immature neutrophils into peripheral tissues.^10,36^ Our GRN modeling results also suggest that tumors need not construct new regulatory pathways to reprogram neutrophils; destabilizing the existing ones is sufficient to lower the barriers that keep homeostatic cells on their normal trajectory and to let them drift into dysfunctional states.^59,60^ More broadly, this type of barrier flattening in the cell state landscape may not be specific to neutrophils, and a comparable loss of landscape structure may occur in other malignancies.^61,62^

Secondly, our perturbation and driving simulations identified *Cebpb* as the dominant driver of the tumor-bearing transition. Driving *Cebpb* alone was sufficient to reproduce the transition while the naive condition instead required two drivers, *Egr1* and *Ets1*, to produce similarly well-fitted results. This is partially explained by the landscape topologies described above. The naive condition is organized into more discrete, barrier-separated states while the tumor-bearing landscape is more continuous and therefore more easily traversed. Biologically, this result is consistent with emergency granulopoiesis, in which *Cebpa* maintains steady-state differentiation checkpoints while *Cebpb* takes over under sustained stress such as the cytokine environment of an expanding tumor.^40,63^ More generally, the fact that a known biological switch emerges as a difference in driving behavior shows NetDes-Duo’s promise for studies in which two conditions can be compared.

Despite these benefits, our approach has several limitations. First, our method is so far limited to modeling only two conditions at a time. Future work may look at extending the method to arbitrarily many conditions as similar metrics to Jaccard overlap can be formulated for this case. Computationally, a more efficient sampling approach would be required as complete enumeration of all combinations of all subsets of regulators would have unacceptably high computational costs with three or more conditions. Second, for the scRNA-seq data, the split into conditions must be supplied in advance. This is applicable to any two-condition scRNA-seq comparison in which both conditions are sampled separately, as in the current neutrophil data. This is generally applicable to comparisons such as treated versus untreated cells, wild type versus knockout, two tissues, or two stages of disease. However, for cases of cell state bifurcation, where two cell fates emerge from a common progenitor, careful preprocessing is required to prepare for two separate datasets for the NetDes-Duo application. Third, our approach does not integrate literature or experimental data to add constraints on condition-specific gene regulation. Doing so could improve the accuracy of the modeling and provide more biologically interpretable networks.

In conclusion, NetDes-Duo provides a general computational approach for modeling the gene regulation of cell state transitions for two related conditions from single-cell gene expression data. By inferring the pair of networks jointly and fitting nonlinear ODEs to each, it recovers structural and dynamical differences. This allows us to turn a comparison of two datasets into two related models that can be simulated, perturbed, and tested for both similarities and differences.

## Methods

### Neutrophil scRNA-seq data and Pre-processing

The scRNA-seq data for neutrophil reprogramming, originally measured in Gong et al. ^36^, were obtained from the Gene Expression Omnibus (dataset GSE217143; samples GSM6705648, GSM6705649, GSM6705650). The six sets of single-cell RNA-sequencing (scRNA-seq) data (neutrophils taken from naive and tumor-bearing mice across three organs: blood, lungs, and bone marrow) were combined into two datasets for downstream processing. These two datasets, consisting of the three naive samples and the three tumor-bearing samples, were designated as the naive and tumor-bearing datasets, respectively. Each dataset underwent independent quality control and pre-processing (log transformation and normalization) in Seurat.^64^ A total of 13,000 high-quality cells were retained in each of the naive and tumor-bearing datasets, roughly evenly divided among the three organ sources. We then conducted principal component analysis (PCA) on the top 500 highly variable genes (HVGs, identified by Seurat’s vst method).^65^

### Smoothed Gene Trajectory Inference

For the dataset of each condition, pseudotime was inferred using a diffusion map via the Destiny package^37^ from the first 20 principal components (PCs). Gene trajectories were smoothed using a LOESS regression (span = 0.75, degree = 2), which applies local quadratic fits to the data. This yielded smoothed gene expression values for each gene at 81 evenly spaced pseudotime points.

### Gene Clustering

Within the naive and tumor-bearing datasets, genes that were both highly expressed and highly variable were selected for clustering. Genes in the top 5000 by variability and above an average-expression cutoff of 0.01 were selected. Approximately 3000 genes fit these criteria.

We then clustered the 3000 smoothed trajectories by their overall trajectory shapes. For each gene, each of the 80 evenly spaced time segments along each of the smoothed trajectories was assigned a label of increasing (i), decreasing (d), or flat (f); after scaling both pseudotime and gene expression to range from 0 to 1, a flat section was considered to be one with an absolute value of slope less than 0.15. Consecutive runs with length *L*>6 of the same label were grouped together, while genes with rare labels were not used for subsequent analyses. This reduction gave each gene a short-ordered sequence of these labels. For example, a label of “ifd” corresponds to a gene expression that initially increases, flattens out, then decreases along its pseudotime trajectory. Genes with the same label were grouped together. This process was carried out for both the naive and tumor-bearing datasets.

This automatic gene grouping scheme generated around 40 trajectory patterns with sizes varying from one to several hundred genes. For each group with size *N* >20, we visually inspected the trajectory density map and manually combined groups with similar dynamics; for example, the groups “d” (decrease) and “df” (decrease-flat) were combined. Following this manual combination, six large gene clusters, each with size *N* >150 remained in both the naive and tumor-bearing datasets.

In the tumor-bearing datasets, the cluster containing the gene group labels “di” and “dfi” (decreasing then increasing) was composed of two clear subsets (**Fig.S2, S3**). In both cases, about half the genes in the cluster reached a minimum one-fourth of the way through pseudotime, and the other half reached a minimum three-fourths of the way through pseudotime. These two subsets were separated, resulting in seven tumor-bearing clusters and six naive clusters, each with size *N* >150 genes.

### Enrichment Analysis

For each dataset, we proceeded to identify core transcription factors (TFs) as components of a gene regulatory network driving observed cell state transitions. We hypothesized that genes with similar expression trajectories were regulated by similar TFs. Therefore, we sought to identify TFs that regulate many genes in the same cluster. To do this, Fisher’s exact test was performed to evaluate the overlap between genes in a cluster and literature-based target genes for a TF, where the background consists of all expressed genes in each respective scRNA-seq dataset. Here, we used a literature-based mouse TF-target database from NetAct^66^. The enrichment analysis resulted in a list of TFs with corresponding *q*-values.

### Obtaining a TF List shared across conditions

Since both datasets were derived from the same biological system, we assumed that their core regulatory networks exhibited high levels of similarity. To reflect this, we took TFs with *q*-value <0.25 in both the naive and tumor-bearing datasets. This resulted in a set of 15 overlapping TFs (*Cebpa*, *Cebpb*, *Ctnnb1*, *Egr1*, *Ets1*, *Ets2*, *Fos*, *Hif1a*, *Id1*, *Jun*, *Mapk13*, *Nfkb2*, *Per1*, *Stat1*, *Stat2*).

Next, we used RcisTarget^18^ to construct TF-TF networks for the naive and tumor-bearing conditions, which applied cis-regulatory motif enrichment to each of the gene clusters using a precomputed motif-gene database. Motifs with Normalized Enrichment Score (NES) >2 were kept. Then, we used TRRUST^35^ to add additional literature-based TF-TF interactions. Finally, we pruned TFs that had either no incoming or no outgoing edges; this corresponded to TFs that were not regulated by, or did not regulate, the rest of the network, respectively. TFs that remained after pruning in both the naive and tumor-bearing cases were kept as the shared core TFs; this yielded 12 shared core TFs (*Cebpa*, *Cebpb*, *Ctnnb1*, *Egr1*, *Ets1*, *Ets2*, *Fos*, *Hif1a*, *Jun*, *Nfkb2*, *Per1*, *Stat1*) compared to the initial 15 overlapping TFs.

### Adding Condition-specific TFs

Using the above-mentioned procedure, we found that, in both the naive and tumor-bearing core TF networks, many TFs were regulated by only one or two other TFs. This is likely because the current networks are missing some critical condition-specific TFs that mediate the network connections. Therefore, we allowed for some additional, non-overlapping TFs to be added to both the naive and tumor-bearing initial networks. For each dataset, we considered the top TFs with the lowest *q*-value that were not already present in the overlapping TF list.

We sought the best subset of these TFs to add to the initial network. For each subset, we first combined the subset with the 12 shared core TFs. Then, we generated an initial network with this combined list using the same process described above (RcisTarget, TRRUST, and pruning). For each subset, we defined a network score as

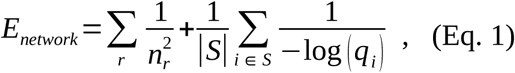

where *n_r_* is the number of regulators for each TF, *S* is the set of additional TFs we added, |*S*| is the size of the set, and *q_i_* is the *q*-value for each of the additional TFs we added. The first term in this score reflected the connectedness of the network, penalizing networks with TFs that only had one or two regulators. The second term in the score reflected the mean *q*-values of the set of additional TFs. We calculated this score for all possible subsets then chose the subset with the lowest score. In the naive case, the TFs *Rel*, *Nfkb1*, and *Stat4* were chosen. In the tumor-bearing case, the TFs *Relb*, *Jund*, and *Sp3* were chosen.

### ODE optimization using NetDes

We next sought to find an optimized network by pruning edges from each initial network. To begin, we utilized NetDes to perform fitting on a nonlinear ODE model. Specifically, NetDes models each TF using a Hill-style ODE given by,

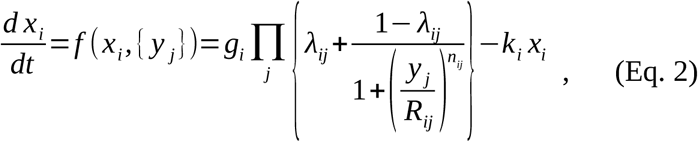

where each multiplicative term represents the transcriptional effect of one regulator. Here *x_i_* is the expression level of the *i*th TF, *y _j_* is the expression level of its *j*th regulator, *g_i_* is the maximum production rate of the *i*th TF, and *k_i_* is the degradation rate. The parameters (*λ_ij_, R_ij_, n_ij_*) are the regulation parameters for the *j*th TF on the *i*th TF. *R_ij_* is the threshold of the Hill function, or the expression level of the *j*th TF at which its regulatory effect on the *i*th TF reaches half of its maximum. *n_ij_* is the Hill coefficient, which controls how steeply the regulation term grows as the regulator crosses *R_ij_*. *λ_ij_* represents the maximum fold change which determines the sign of the regulation, with *λ_ij_*>1 meaning activation and *λ_ij_* <1 meaning repression. During fitting each *λ* is constrained between 0.01< *λ*< 100.

NetDes performs parameter fitting to minimize mean squared error (MSE). NetDes first samples many initial guesses to avoid poor local minima and move toward a global optimum. Top initial guesses go through two rounds of refinement, generating new starting points based on the top successful guesses from previous rounds, and a five-fold cross-validation (CV) process. This yields a list of the 10 models with the lowest MSE.

Next, we sought to retain only core regulation terms. For a given TF and a chosen subset of its regulators, we constrained the model to keep only that subset of regulators, whereas all other regulators were deleted from the corresponding ODE. We performed one round of re-optimization on the remaining parameters, initializing from the previously fitted parameter values. For each possible subset of regulators, we obtained a list of the 10 models with the lowest MSE. We then compared these against the list of the top 10 models before deletion. Specifically, NetDes has previously defined a reducibility index,

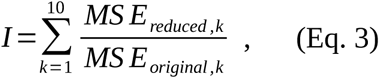

A reducibility value closer to 10 means that the subset performs nearly as well as the full set of regulators, while a higher value suggests a key regulator has been lost.

### Combined Deletion to Maximize Overlap

Many, but not all, of the pathways and core TF-TF interactions remain unchanged between the two conditions. Therefore, we sought to obtain a pair of optimized networks that are highly overlapping. To this end, we sought a deletion metric that prioritizes deleting disagreeing TF-TF connections. At the same time, both subsets should still have low reducibility indices.

To accomplish this, we defined a combined scoring metric that utilizes both Jaccard index and the reducibility index from both conditions. For a given shared TF, suppose it is regulated by a set of regulators *R_N_* in the initial network of the first condition and a set of regulators *R_C_* in the initial network of the second condition. To begin, we calculated the reducibility indices for all subsets of *R_N_* and for all subsets of *R_C_* . Then, across all pairs of subsets from the two conditions, we calculated,

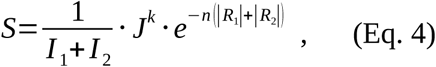

where *R*_1_ and *R*_2_ are the subsets of regulators; |*R*_1_| and |*R*_2_| are the sizes of those subsets; *I* _1_ and *I* _2_ are their corresponding reducibility indices; and *J* is the Jaccard index defined as

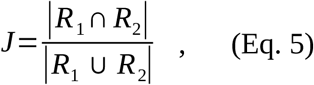

In particular, the Jaccard index is calculated such that *R*_1_ *∩ R*_2_ only includes connections that agree in sign (i.e., they must both be activating interactions or both be repressing interactions). Specifically, we only count a given interaction (*λ_ij_*, see Eq. 2) as part of the intersection if the two models with lowest MSE have either *λ_ij, N_, λ_ij,C_* <1 or *λ_ij, N_, λ_ij,C_* >1. The two subsets in the pair with maximum score were chosen for the corresponding optimized networks.

If we were dealing with a non-overlapping TF that did not exist in the other network, we instead used the following score to find the subset of regulators to keep,

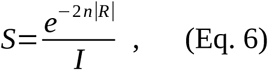

where *I* is the corresponding reducibility index to *R*_2_. This expression was obtained by considering the case of *R*_1_=*R*_2_ from the previous scoring metric. Once again, the subset with maximum score was chosen for the optimized network.

This combined score simultaneously rewards low reducibility indices, high overlap between regulator subsets, and smaller overall regulator sets. Here, we have two hyperparameters *n* and *k*, which can be used to tune the overall network; larger *n* values lead to a smaller overall network, and larger *k* values lead to a higher overlap percentage between the networks.

### Hyperparameter Selection

To choose appropriate parameters, we ran the entire NetDes-Duo optimization pipeline across different values of *n*, resulting in many final optimized ODE models. We plotted the average MSE of these fits against *n* (**Fig.S6A**), while holding *k* constant at 1. We noticed a steep increase in the tumor-bearing network beyond *n*=0.5, corresponding to a network size around 40 (**Fig.S6B**). Therefore, we chose *n*=0.5 for both networks so that they would have similar size. To choose an appropriate *k*, we plotted the average MSE increase of the fitting against *k,* while holding *n*=1 (**Fig.S6C**). Here, *k* =1.5 was selected as it produced a relatively low average MSE in both networks.

## Landscape Characterization

Using the optimized ODE models, we characterized the cell state behavior with model simulations. To do this, we randomly sampled 5000 initial states, where each gene expression was chosen randomly and independently between the minimum and maximum experimental levels. To map the regulatory landscape, we performed stochastic simulations by adding noise terms to the drift terms derived from the optimized ODE. Specifically, for each gene *i*

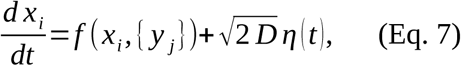

where *f* is the deterministic term defined in Eq. 2, *D*=8*×* 10^−4^ sets the noise strength corresponding to a noise amplitude _√_2 *D* = 0.04, and *η* (*t*) is Gaussian white noise, uncorrelated in time and independent across genes. The expression level of every gene was scaled to the interval [0,1], so the same noise strength was applied to all genes. For each of the 5000 cases, we numerically integrated the stochastic differential equations (3000 time steps, step size 1) resulting in gene expression profiles of 5000 simulated cells.

Next, we applied PCA on these simulated cells and applied the R package princurve^67^ to obtain a principal curve through the cloud. Finally, we calculated quasi-potential along this principal curve. Specifically, we estimated the probability for a state to be at any given position along the principal curve via kernel density estimation. Then, we defined quasi-potential as

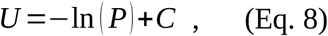

where *C* is a constant chosen per condition so that the global minimum of each quasi-potential is zero.

As a check on our model, we projected the naive and tumor-bearing single-cell neutrophil data onto the derived PCA. For each condition, the log expression for each of the 15 TFs used was standardized to zero mean and unit variance. The standardized vectors from the naive and tumor-bearing data were then projected onto their respective simulated PCAs. Then, each cell was assigned a principal curve coordinate by orthogonal projection onto its respective principal curve. Because the single-cell data are noisy, we performed this projection with a stretch parameter to allow some measured cells to fall beyond the 0 to 1 range that all simulated cells are within. Finally, we confirmed the relative ordering of cells along the simulated principal curve by visually inspecting the histograms of cell states by tissue type (**Fig.5C**).

### Perturbation simulations

To identify key transcription factors that can drive the cell-state transition, we performed perturbation simulations by transiently over-expressing specific gene(s). We began by sampling final states from the 5000 simulated cells produced by the unperturbed stochastic simulation. In this way, each perturbation is applied across the full baseline distribution of states rather than to cells from a single basin. From here, the expression of a single gene or pair of genes was increased by

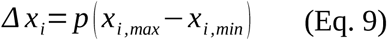

very quickly over the course of some small number of time steps (here we chose 20). Here, *p* is a coefficient that determines the strength of the perturbation. Then, the system was allowed to relax by a stochastic simulation that integrates Eq. 7 for another 2000 steps. Because the perturbation is applied at a single time point and the underlying differential equation models are unchanged, we expect that any perturbation will fade away with enough time. In this case, we observed large shifts after 2000 steps, so the time scale for full decay is likely much larger; the 2,000-step window captures displacements that persist over finite times rather than changes in the equilibrium distribution.

After the relaxation period, the resulting simulated states were projected onto the first two principal components and principal curve derived from the corresponding condition’s baseline cloud, assigning each perturbed state a position along the same principal curve used to characterize the unperturbed landscape. We used the average shift in curve position to quantify the strength of the perturbation. This projection was performed separately for the naive and tumor-bearing conditions across all 78 pairs and all 12 single TFs drawn from the core TF set. For a few top perturbations, we also varied the size of the perturbation (*p* from above) and recorded the response in the curve-position shift.

### Driving simulations

As a second approach to identify TFs responsible for the observed cell-state transitions, we performed driving simulations. We took the starting point from each gene’s smoothed time trajectories and simulated the optimized ODEs until a steady state was reached; by the nature of the ODE fitting, this steady state was very close to each of the gene’s starting points. Next, for either a single TF or a pair of TFs, we clamped the expression of the driven TF(s) to their fitted trajectories from the scRNA-seq data. We first performed this simulation forward in time. To check for hysteresis, we then used the final state of the forward simulation as a new starting point and drove the system backward using the same clamped gene expression trajectory/trajectories in reverse.

### Synthetic Data Generation for Benchmarking

To benchmark NetDes-Duo with existing network inference methods, we generated pairs of highly similar gene regulatory networks. For both networks, the gene expression dynamics of one driving gene were fixed with a formula,

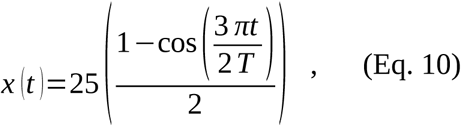

where *T* is our total simulation time. We then randomly sampled network structures with a fixed number of edges, subject to the constraint that each gene must have at least one incoming and one outgoing edge. For each edge, the parameter set (*λ_ij_, R_ij_, n_ij_*) from Eq. 2 was randomly sampled where log (*λ_ij_*) ranged from −log (6) to −log (3) for inhibition and log (6) to log (3) for activation; *R_ij_* was sampled from 0.1 to 0.5 of the gene expression range; and log (*n_ij_*) was sampled from log (1) to log (5).

To generate the second network in each pair, we randomly sampled network structures with the same fixed number of edges, where we additionally applied the constraint that the new edge set must have Jaccard similarity equal to or greater than 0.7 with that of the first network. For overlapping edges, we also ensured the sign (activation/inhibition) was consistent between networks. The other parameters, along with the specific value of *λ_ij_*, were all re-sampled. Thus, the pair of networks had similar but non-identical structures, and often substantially different gene trajectories.

Most randomly sampled network pairs produced groups of 3-4 highly correlated genes with the Pearson correlation coefficient often being >0.995 between pairs of genes. Downstream analysis with these pairs proved difficult: neither our ODE model nor the other benchmarking methods we used could distinguish between these very similar genes. To avoid this issue, we computed the maximum Pearson correlation coefficient across all pairs of gene trajectories for each network. We set a similarity cutoff at the bottom 1% of these maximum-similarity values. In addition, for each pair of networks, we recorded the maximum slope of the normalized gene expression across all genes. We set a cutoff at the bottom 80% of maximum slopes to filter out networks with oscillatory behavior. 2000 pairs of networks were generated to obtain numerical values for these cutoffs. Then, we continued to generate pairs of networks and retained only those that passed both the similarity and slope cutoffs. Approximately 6000 pairs were generated to obtain 20 pairs that passed both cutoffs.

To test whether our method recovers the correct edge set, we added varying numbers of decoys to each initial network. We added between 9 and 24 decoy edges to a network of 18 ground-truth edges and 7 genes, resulting in initial networks with 27-42 total edges.

### Comparison to Other Regulatory Network Inference Methods

We employed multiple regulatory network reconstruction methods, including GENIE3^68^, ppcor^69^, SCODE^70^, DeepSEM^22^, and NetDes^29^, alongside NetDes-Duo. GENIE3 and DeepSEM were used for unsigned interaction recovery, while ppcor, SCODE, and NetDes were additionally used for signed interaction comparisons. All methods were run on default settings, although the size of each network was matched to that of NetDes-Duo. The implementation of these methods is outlined as follows.

The ppcor method calculates partial and semi-partial correlation coefficients between candidate regulators and target genes. We took the semi-partial correlation as the interaction score and used its sign to label each interaction as activating or inhibiting. Candidate regulatory interactions were ranked by the absolute value of the semi-partial correlation coefficient.

GENIE3 employs a tree-based regression model to predict regulatory interactions between genes. For each target gene it fits a tree ensemble predicting that gene’s expression from the expression of all candidate regulators and takes each regulator’s importance in that model as the edge weight. We applied GENIE3 to the processed synthetic data using its default settings. Regulatory edges were ranked by the interaction weights and were evaluated only for unsigned recovery.

SCODE infers regulatory networks by fitting gene-expression dynamics with a linear ODE. Each inferred coefficient was labeled activating or inhibiting according to the sign of the coefficient, and edges were ranked by the magnitude of the coefficient. We ran 100 iterations of optimization on default settings.

DeepSEM learns a latent representation of the processed expression matrix with a neural network and derives gene-gene regulatory weights from it. We applied DeepSEM to the synthetic data using its default settings. The inferred edge weights were used to rank candidate regulatory interactions, and DeepSEM was evaluated only for unsigned recovery of regulator-target edges.

NetDes’s original deletion algorithm uses the same reducibility index *I* as defined above. However, in order to achieve a reduced network, it utilizes a cutoff *C*, and selects the smallest possible subset of regulators that satisfies *I* < *C* for each TF. In order to generate a variety of network sizes, we swept many cutoff values ranging from 10 to 1000.

### Evaluating Performance of Methods on Synthetic Data

For NetDes-Duo, we fixed *k* at 1 and swept *n* from 0 to 100, yielding inferred networks that span a wide range of sizes. We applied all the above methods to all 40 networks separately, while NetDes-Duo was applied to the 20 associated pairs of networks. To evaluate the performance of each method, we evaluated AUPRC on the whole network. Specifically, at each network size, we calculated the precision and recall of the inferred edges across the whole network relative to the ground truth network and then integrated the resulting precision-recall curve to obtain the area. We also calculated an activating and inhibiting interaction AUPRC for methods that assign a sign to each interaction. Here, the methods must not only identify the decoys successfully but also distinguish between activating and inhibiting edges. Specifically, an activating edge being labeled as an inhibiting edge is counted as a false negative for the activating AUPRC and a false positive for the inhibiting AUPRC.

## Code availability

The implementation of NetDes-Duo along with all scripts required to reproduce the synthetic analysis, the neutrophil analysis, and the figures in this paper are available at https://github.com/lusystemsbio/NetDes-Duo.

## Data availability

The scRNA-seq data analyzed in this study are publicly available from the Gene Expression Omnibus under accession GSE217143 (samples GSM6705648, GSM6705649, and GSM6705650). The synthetic benchmarking data along with fitted networks are available at https://github.com/lusystemsbio/NetDes-Duo.

## Acknowledgement

We thank the Bioengineering Department at Northeastern University and its minor programs for hosting A. Ren’s research internship in the Lu lab. A. Ren, Y. You, and M. Lu are supported by startup funds from Northeastern University. Y. You and M. Lu are also supported by the National Institute of General Medical Sciences of the National Institutes of Health under Award Number R35GM128717. A. Ren, Y. You, and M. Lu acknowledge their affiliation with the Center for Theoretical Biological Physics at Northeastern University and appreciate the support provided by the center.

## AI Statement

The authors used Claude to check grammar and improve clarity of the manuscript text. The authors reviewed and edited all AI-assisted output and take full responsibility for the content of the manuscript.

## References

1. Wang, J., Zhang, K., Xu, L. & Wang, E. Quantifying the Waddington landscape and biological paths for development and differentiation. Proc. Natl. Acad. Sci. U. S. A. 108, 8257–8262 (2011).

2. Marjanovic, N. D. et al. Emergence of a High-Plasticity Cell State during Lung Cancer Evolution. Cancer Cell 38, 229–246.e13 (2020).

3. Huang, S., Guo, Y.-P., May, G. & Enver, T. Bifurcation dynamics in lineage-commitment in bipotent progenitor cells. Dev. Biol. 305, 695–713 (2007).

4. Costa, A. et al. Fibroblast Heterogeneity and Immunosuppressive Environment in Human Breast Cancer. Cancer Cell 33, 463–479.e10 (2018).

5. Confalonieri, P. et al. Regeneration or Repair? The Role of Alveolar Epithelial Cells in the Pathogenesis of Idiopathic Pulmonary Fibrosis (IPF). Cells 11, 2095 (2022).

6. Ng, M., Cerezo-Wallis, D., Ng, L. G. & Hidalgo, A. Adaptations of neutrophils in cancer. Immunity 58, 40–58 (2025).

7. Ballesteros, I. et al. Co-option of Neutrophil Fates by Tissue Environments. Cell 183, 1282–1297.e18 (2020).

8. Evrard, M. et al. Developmental Analysis of Bone Marrow Neutrophils Reveals Populations Specialized in Expansion, Trafficking, and Effector Functions. Immunity 48, 364–379.e8 (2018).

9. Kwok, I. et al. Combinatorial Single-Cell Analyses of Granulocyte-Monocyte Progenitor Heterogeneity Reveals an Early Uni-potent Neutrophil Progenitor. Immunity 53, 303–318.e5 (2020).

10. Eruslanov, E., Nefedova, Y. & Gabrilovich, D. I. The heterogeneity of neutrophils in cancer and its implication for therapeutic targeting. Nat. Immunol. 26, 17–28 (2025).

11. Casbon, A.-J. et al. Invasive breast cancer reprograms early myeloid differentiation in the bone marrow to generate immunosuppressive neutrophils. Proc. Natl. Acad. Sci. U. S. A. 112, E566–575 (2015).

12. Haghverdi, L. & Ludwig, L. S. Single-cell multi-omics and lineage tracing to dissect cell fate decision-making. Stem Cell Rep. 18, 13–25 (2023).

13. Wang, W. et al. Live-cell imaging and analysis reveal cell phenotypic transition dynamics inherently missing in snapshot data. Sci. Adv. 6, eaba9319 (2020).

14. Wu, X. et al. Single-cell sequencing to multi-omics: technologies and applications. Biomark. Res. 12, 110 (2024).

15. Saelens, W., Cannoodt, R., Todorov, H. & Saeys, Y. A comparison of single-cell trajectory inference methods. Nat. Biotechnol. 37, 547–554 (2019).

16. Schiebinger, G. et al. Optimal-Transport Analysis of Single-Cell Gene Expression Identifies Developmental Trajectories in Reprogramming. Cell 176, 928–943.e22 (2019).

17. Chan, T. E., Stumpf, M. P. H. & Babtie, A. C. Gene Regulatory Network Inference from Single-Cell Data Using Multivariate Information Measures. Cell Syst. 5, 251–267.e3 (2017).

18. Aibar, S. et al. SCENIC: single-cell regulatory network inference and clustering. Nat. Methods 14, 1083–1086 (2017).

19. Papili Gao, N., Ud-Dean, S. M. M., Gandrillon, O. & Gunawan, R. SINCERITIES: inferring gene regulatory networks from time-stamped single cell transcriptional expression profiles. Bioinformatics 34, 258–266 (2018).

20. Wang, L. et al. Dictys: dynamic gene regulatory network dissects developmental continuum with single-cell multiomics. Nat. Methods 20, 1368–1378 (2023).

21. Zhang, S. et al. Inference of cell type-specific gene regulatory networks on cell lineages from single cell omic datasets. Nat. Commun. 14, 3064 (2023).

22. Shu, H. et al. Modeling gene regulatory networks using neural network architectures. *Nat. Comput. Sci.* 1, 491–501 (2021).

23. Yuan, Q. & Duren, Z. Inferring gene regulatory networks from single-cell multiome data using atlas-scale external data. Nat. Biotechnol. 43, 247–257 (2025).

24. 24. Bertin, P., et al. A scalable gene network model of regulatory dynamics in single cells. Preprint at 10.48550/ARXIV.2503.20027 (2025).

25. Qiu, X. et al. Mapping transcriptomic vector fields of single cells. Cell 185, 690–711.e45 (2022).

26. Kamimoto, K. et al. Dissecting cell identity via network inference and in silico gene perturbation. Nature 614, 742–751 (2023).

27. Wang, W. et al. RegVelo: Gene-regulatory-informed dynamics of single cells. Cell 189, 3773–3800.e44 (2026).

28. Chen, F. & Li, C. Reconstructing gene network structure and dynamics from single cell data. Bioinformatics 41, btaf598 (2025).

29. You, Y., Caranica, C. & Lu, M. Building dynamical models of multi-step state transitions from single cell gene expression trajectories. Preprint at 10.64898/2025.12.08.693064 (2025).

30. Katebi, A., Ramirez, D. & Lu, M. Computational systems-biology approaches for modeling gene networks driving epithelial–mesenchymal transitions. Comput. Syst. Oncol. 1, e1021 (2021).

31. Jia, C. et al. Accounting for technical noise in differential expression analysis of single-cell RNA sequencing data. Nucleic Acids Res. 45, 10978–10988 (2017).

32. Pratapa, A., Jalihal, A. P., Law, J. N., Bharadwaj, A. & Murali, T. M. Benchmarking algorithms for gene regulatory network inference from single-cell transcriptomic data. Nat. Methods 17, 147–154 (2020).

33. Zhou, X. & Cai, X. Inference of differential gene regulatory networks based on gene expression and genetic perturbation data. Bioinformatics 36, 197–204 (2020).

34. Duren, Z. et al. Sc-compReg enables the comparison of gene regulatory networks between conditions using single-cell data. Nat. Commun. 12, 4763 (2021).

35. Han, H. et al. TRRUST v2: an expanded reference database of human and mouse transcriptional regulatory interactions. Nucleic Acids Res. 46, D380–D386 (2018).

36. Gong, Z., et al. Immunosuppressive reprogramming of neutrophils by lung mesenchymal cells promotes breast cancer metastasis. Sci. Immunol. 8, eadd5204 (2023).

37. Angerer, P., et al. *destiny* : diffusion maps for large-scale single-cell data in R. Bioinformatics 32, 1241–1243 (2016).

38. Zhang, D. E. et al. Absence of granulocyte colony-stimulating factor signaling and neutrophil development in CCAAT enhancer binding protein alpha-deficient mice. Proc. Natl. Acad. Sci. U. S. A. 94, 569–574 (1997).

39. Avellino, R. et al. An autonomous CEBPA enhancer specific for myeloid-lineage priming and neutrophilic differentiation. Blood 127, 2991–3003 (2016).

40. Hirai, H. et al. C/EBPbeta is required for ‘emergency’ granulopoiesis. Nat. Immunol. 7, 732–739 (2006).

41. Satake, S. et al. C/EBPβ is involved in the amplification of early granulocyte precursors during candidemia-induced ‘emergency’ granulopoiesis. J. Immunol. 189, 4546–4555 (2012).

42. Atsaves, V., Leventaki, V., Rassidakis, G. Z. & Claret, F. X. AP-1 Transcription Factors as Regulators of Immune Responses in Cancer. Cancers 11, 1037 (2019).

43. Khoyratty, T. E. et al. Distinct transcription factor networks control neutrophil-driven inflammation. Nat. Immunol. 22, 1093–1106 (2021).

44. Walmsley, S. R. et al. Hypoxia-induced neutrophil survival is mediated by HIF-1alpha-dependent NF-kappaB activity. J. Exp. Med. 201, 105–115 (2005).

45. Ovadia, S., Özcan, A. & Hidalgo, A. The circadian neutrophil, inside-out. J. Leukoc. Biol. 113, 555–566 (2023).

46. Adrover, J. M. et al. A Neutrophil Timer Coordinates Immune Defense and Vascular Protection. Immunity 50, 390–402.e10 (2019).

47. Wang, H., Zhou, K., Li, W., Du, J. & Xiao, J. Ctnnb1 transcriptional upregulation compensates for Mdm2/p53-mediated β-catenin degradation in neutrophils following cardioembolic stroke. Gene 766, 145022 (2021).

48. Cullen, E. M., Brazil, J. C. & O’Connor, C. M. Mature human neutrophils constitutively express the transcription factor EGR-1. Mol. Immunol. 47, 1701–1709 (2010).

49. Sun, S.-C. The non-canonical NF-κB pathway in immunity and inflammation. Nat. Rev. Immunol. 17, 545–558 (2017).

50. Yoshida, S. et al. Interferon-γ induces interleukin-6 production by neutrophils via the Janus kinase (JAK)-signal transducer and activator of transcription (STAT) pathway. BMC Res. Notes 14, 447 (2021).

51. Xiao, G., Harhaj, E. W. & Sun, S. C. NF-kappaB-inducing kinase regulates the processing of NF-kappaB2 p100. Mol. Cell 7, 401–409 (2001).

52. Li, F. et al. Single-cell atlas reveals a pro-metastatic RELB+ neutrophil-myeloid subset underlying lymph node metastasis in EGFR-wildtype LUAD. Front. Cell Dev. Biol. 14, 1766211 (2026).

53. Duveau, C. et al. Key role of transcription factors network in proliferative vitreoretinal diseases development. Cell Biosci. 16, 71 (2026).

54. Gào, X. & Schöttker, B. Reduction-oxidation pathways involved in cancer development: a systematic review of literature reviews. Oncotarget 8, 51888–51906 (2017).

55. Marigo, I. et al. Tumor-induced tolerance and immune suppression depend on the C/EBPbeta transcription factor. Immunity 32, 790–802 (2010).

56. Saadeddin, A., Babaei-Jadidi, R., Spencer-Dene, B. & Nateri, A. S. The links between transcription, beta-catenin/JNK signaling, and carcinogenesis. Mol. Cancer Res. MCR 7, 1189–1196 (2009).

57. Hidalgo, A., Chilvers, E. R., Summers, C. & Koenderman, L. The Neutrophil Life Cycle. Trends Immunol. 40, 584–597 (2019).

58. Overbeeke, C., Tak, T. & Koenderman, L. The journey of neutropoiesis: how complex landscapes in bone marrow guide continuous neutrophil lineage determination. Blood 139, 2285–2293 (2022).

59. de Visser, K. E. & Joyce, J. A. The evolving tumor microenvironment: From cancer initiation to metastatic outgrowth. Cancer Cell 41, 374–403 (2023).

60. Savy, T., Flanders, L., Karpanasamy, T., Sun, M. & Gerlinger, M. Cancer evolution: from Darwin to the Extended Evolutionary Synthesis. Trends Cancer 11, 204–215 (2025).

61. Burkhardt, D. B., San Juan, B. P., Lock, J. G., Krishnaswamy, S. & Chaffer, C. L. Mapping Phenotypic Plasticity upon the Cancer Cell State Landscape Using Manifold Learning. Cancer Discov. 12, 1847–1859 (2022).

62. Pérez-González, A., Bévant, K. & Blanpain, C. Cancer cell plasticity during tumor progression, metastasis and response to therapy. *Nat*. Cancer 4, 1063–1082 (2023).

63. Manz, M. G. & Boettcher, S. Emergency granulopoiesis. Nat. Rev. Immunol. 14, 302–314 (2014).

64. Hao, Y. et al. Integrated analysis of multimodal single-cell data. Cell 184, 3573–3587.e29 (2021).

65. Hao, Y. et al. Dictionary learning for integrative, multimodal and scalable single-cell analysis. Nat. Biotechnol. 42, 293–304 (2024).

66. Su, K. et al. NetAct: a computational platform to construct core transcription factor regulatory networks using gene activity. Genome Biol. 23, 270 (2022).

67. Cannoodt, R. & Bengtsson, H. rcannood/princurve: princurve 2.1.4. Zenodo 10.5281/ZENODO.3351282 (2019).

68. Huynh-Thu, V. A., Irrthum, A., Wehenkel, L. & Geurts, P. Inferring Regulatory Networks from Expression Data Using Tree-Based Methods. PLoS ONE 5, e12776 (2010).

69. Kim, S. ppcor: An R Package for a Fast Calculation to Semi-partial Correlation Coefficients. Commun. Stat. Appl. Methods 22, 665–674 (2015).

70. Matsumoto, H. et al. SCODE: an efficient regulatory network inference algorithm from single-cell RNA-Seq during differentiation. Bioinformatics 33, 2314–2321 (2017).

