## Supplemental Figures for "Joint inference of paired dynamical gene regulatory networks reveals distinct cell-state landscapes of neutrophil reprogramming"

A

### Networks 1-20 (pairs 1-10)

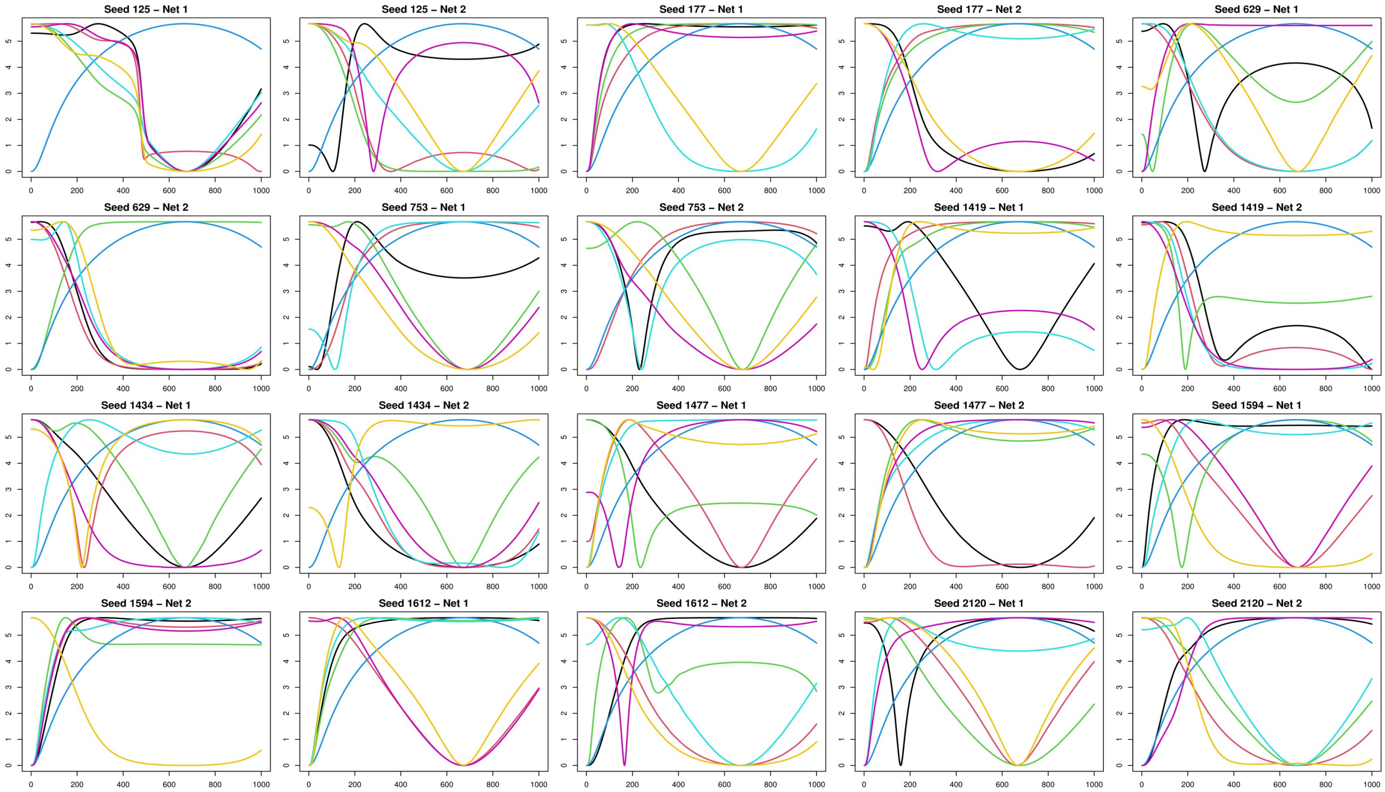

B

#### Networks 21-40 (pairs 11-20)

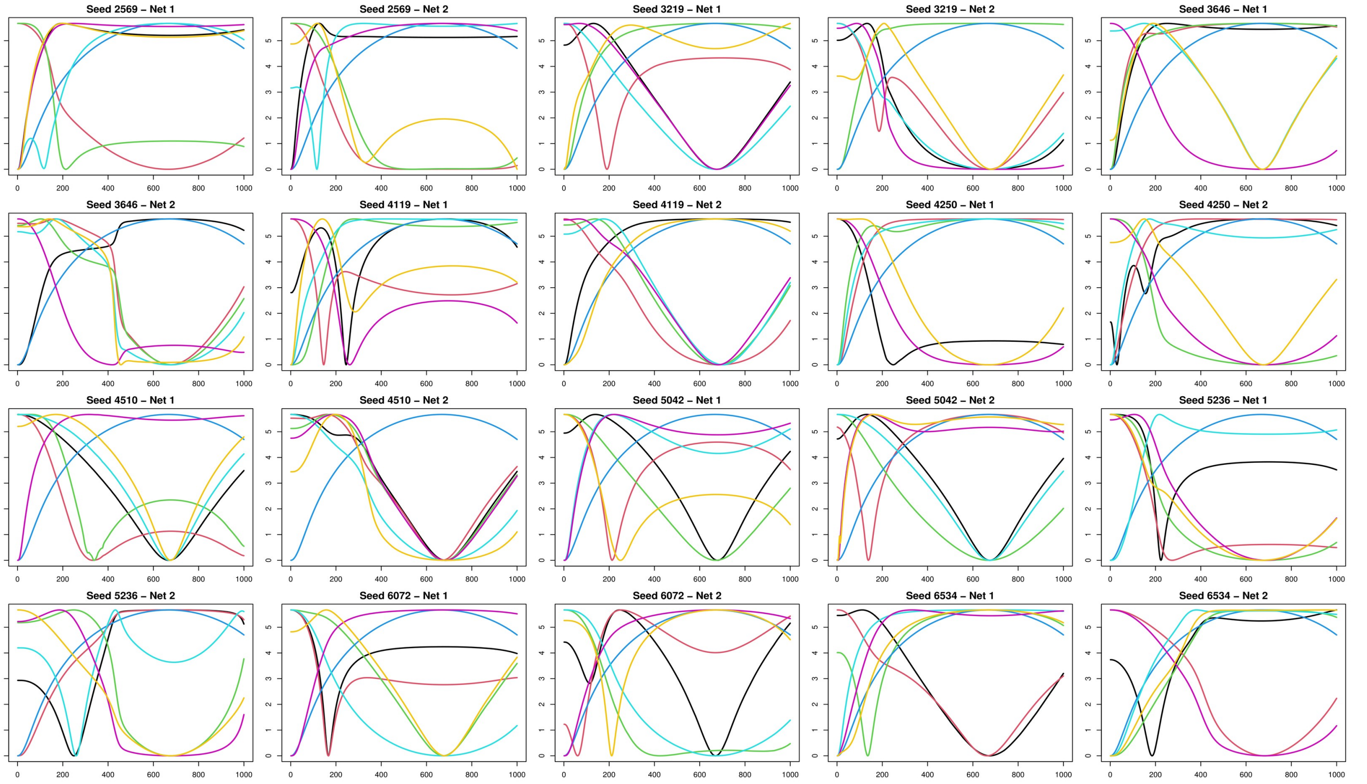


**Fig S1. Simulated gene expression trajectories for all synthetic benchmark networks. (A)** Trajectories for networks 1-20. **(B)** Trajectories for networks 21-40. Paired networks share at least 70% of their edges by Jaccard index but have independently sampled kinetic parameters; one representative pair is shown in **Fig.2A**.

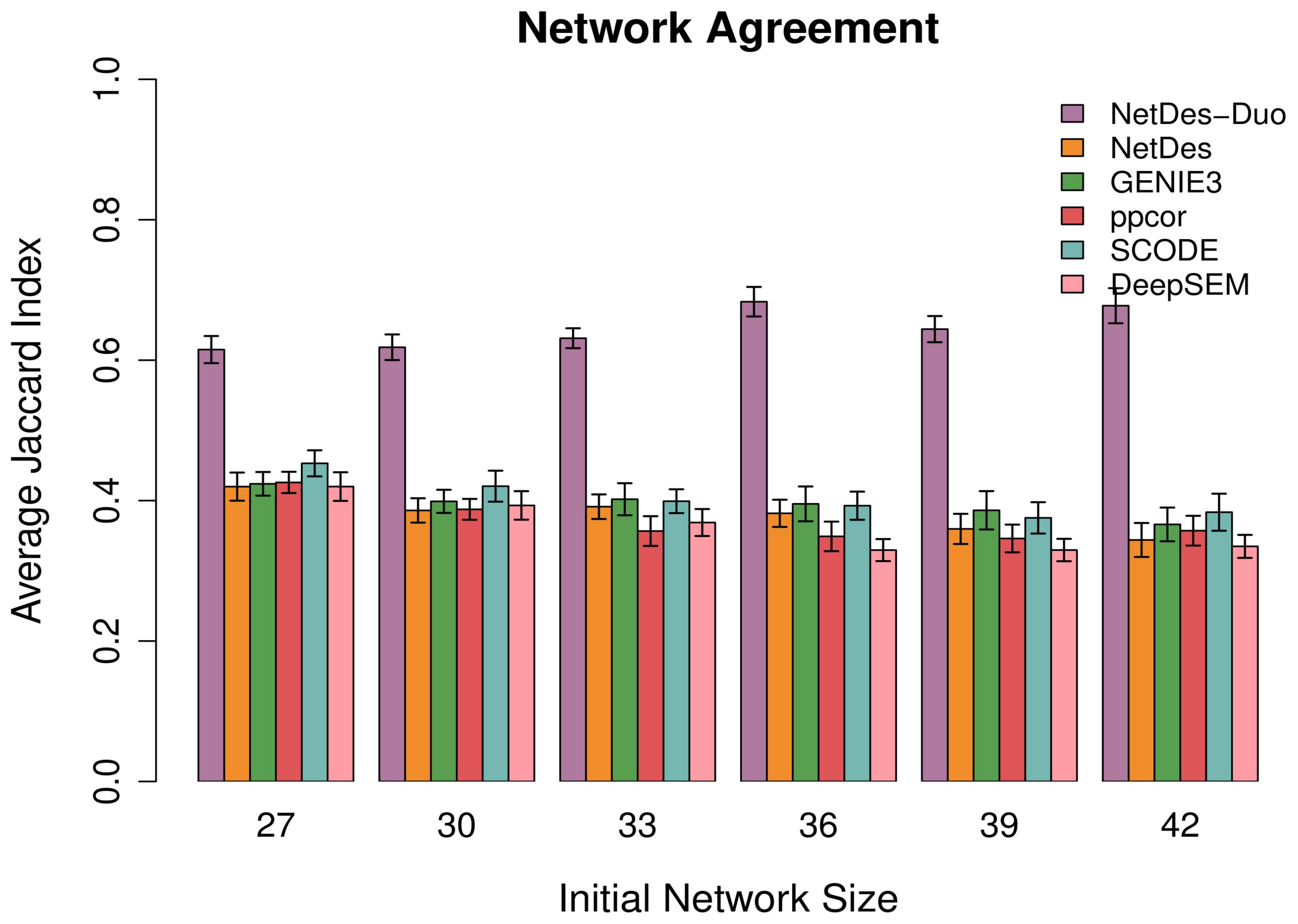


**Fig S2. Agreement between synthetic networks by method.** Average Jaccard index between the two inferred networks of each benchmark pair, for NetDes-Duo and the five comparison methods, across a range of initial network sizes. Bars show the mean across all 20 benchmark pairs and error bars the standard error.

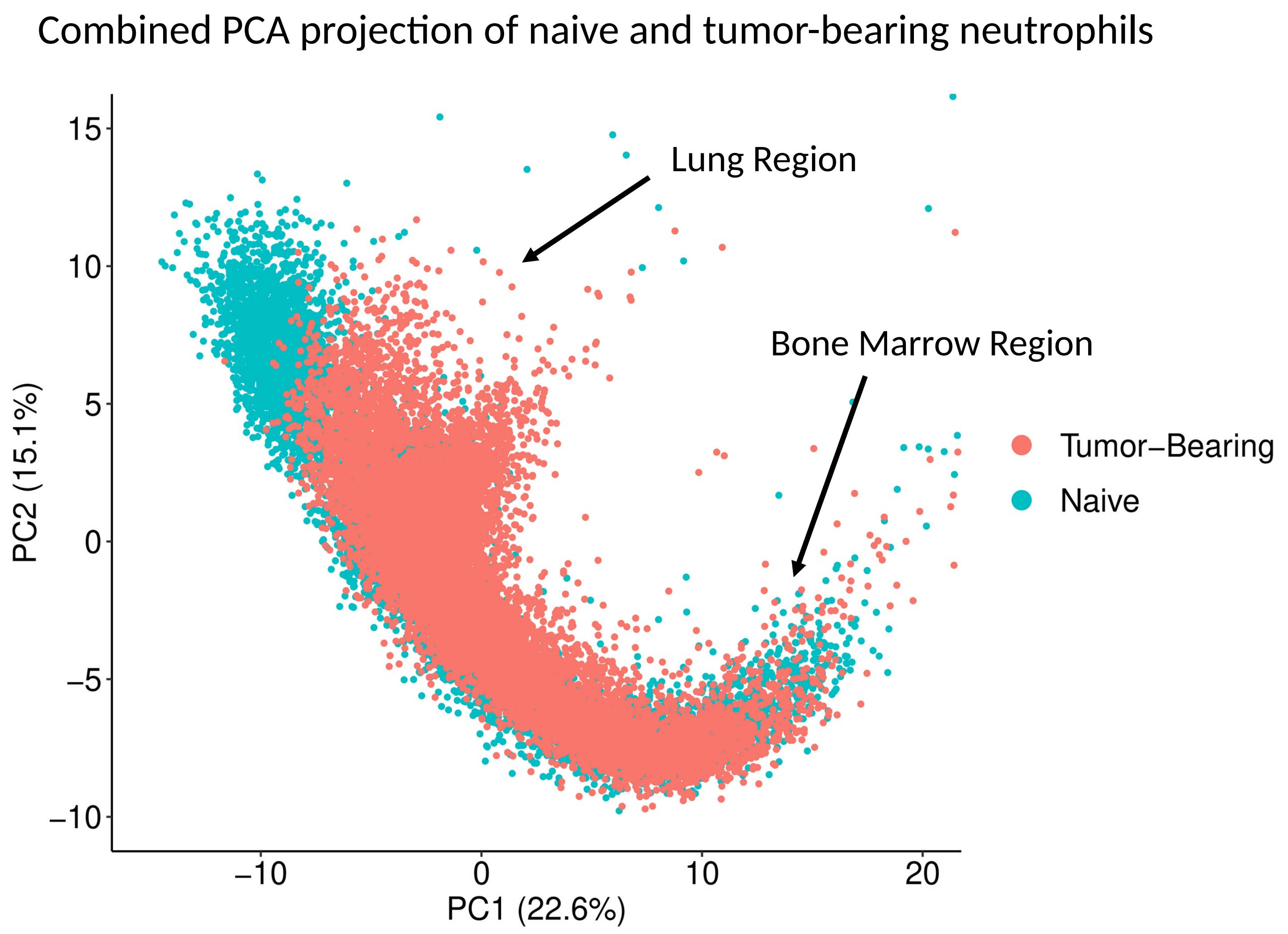


**Fig S3. Combined PCA projection of naive and tumor-bearing neutrophils.** Naive and tumor-bearing single-cell profiles projected together onto the first two principal components. The two conditions overlap in the bone marrow region and diverge toward distinct lung-associated states.

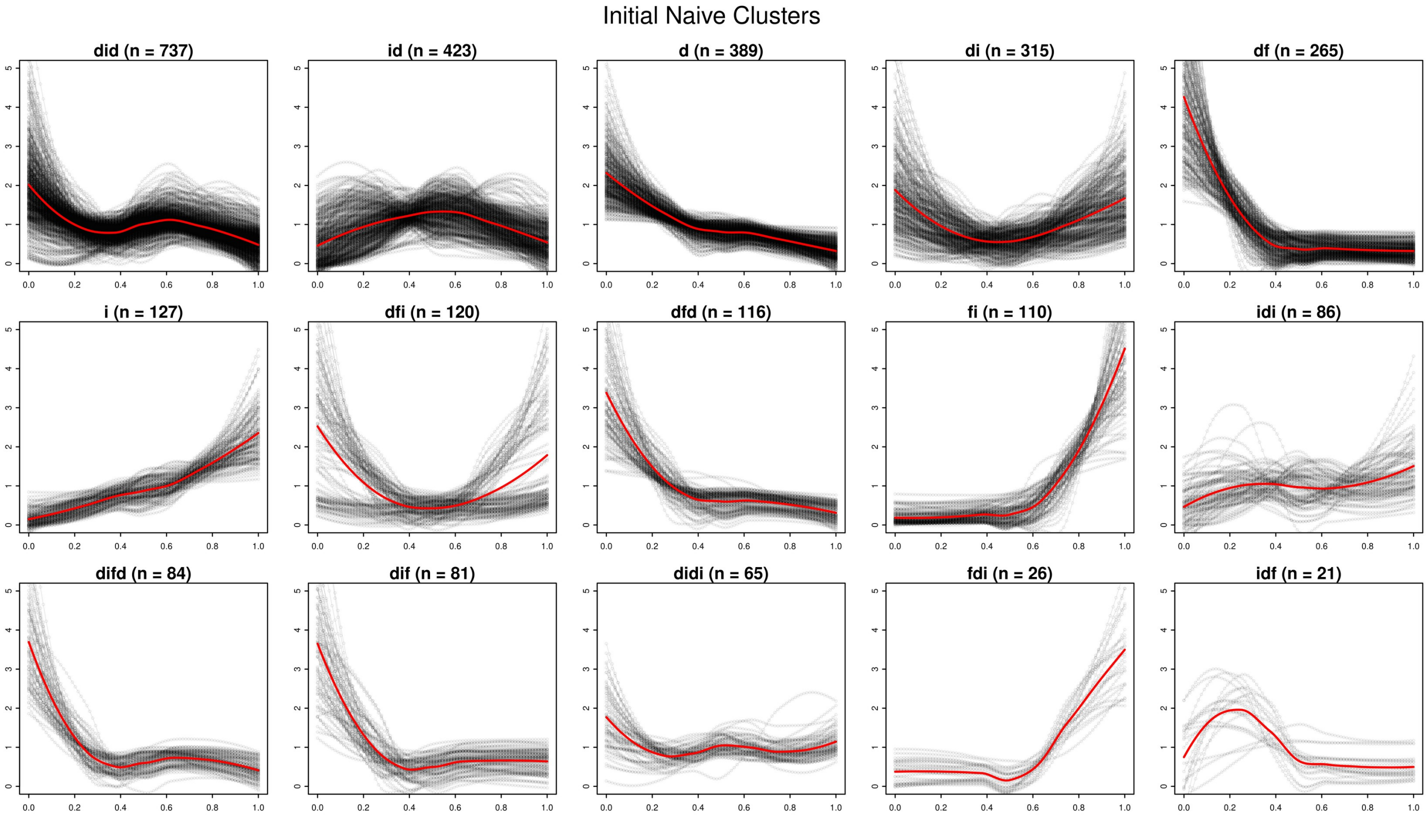


**Fig S4. Automatic trajectory clustering of the naive dataset.** Each panel is one automatically generated gene group for the naive condition, labeled by its sequence of increasing (i), decreasing (d), and flat (f) segments. Grey curves are individual smoothed gene trajectories, and the red curve is the group mean. Groups with N > 20 were combined manually into the six naive clusters shown in **Fig.3B**, with the “d” and “di” groups being further split into two.

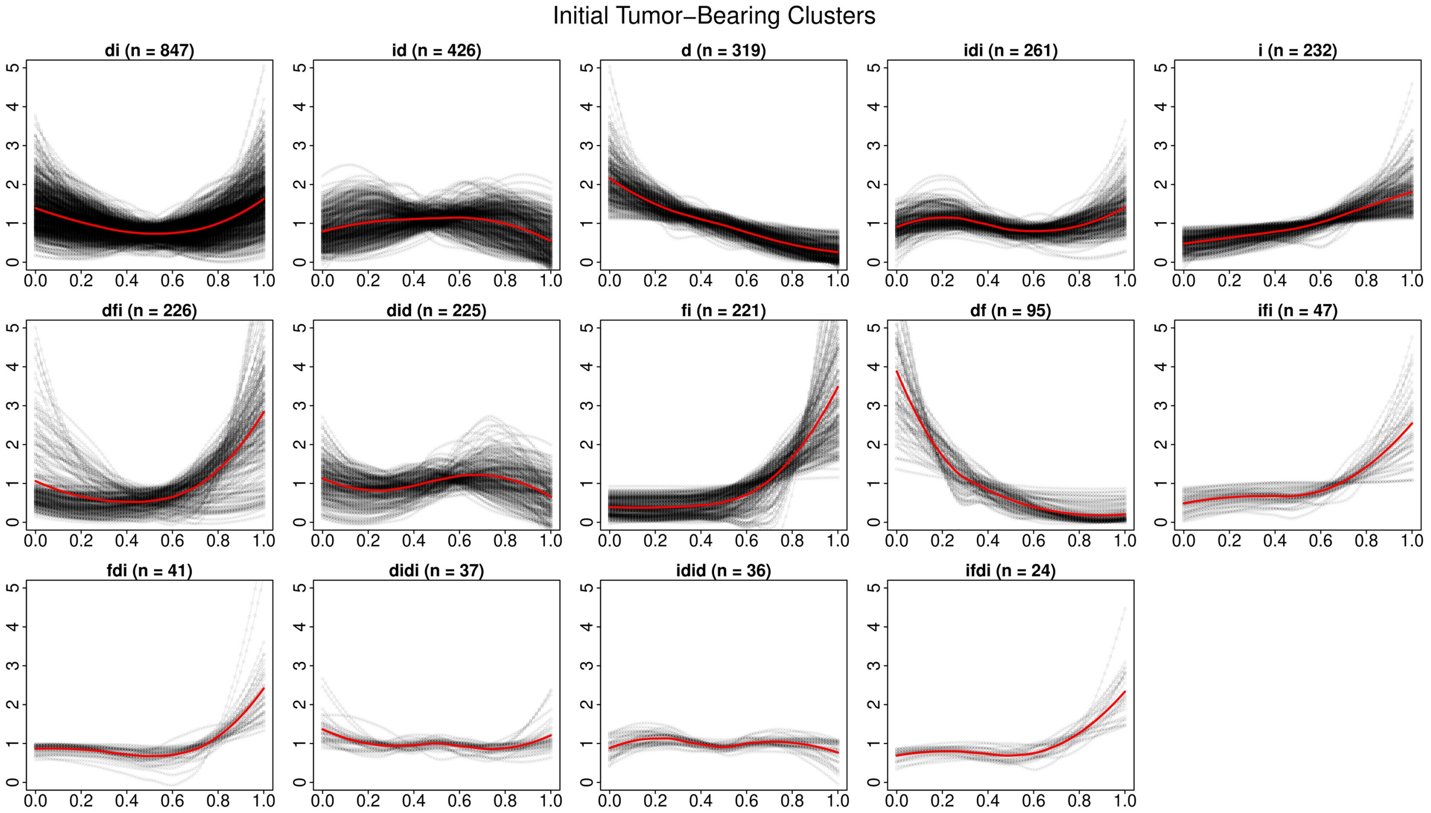


**Fig S5. Automatic trajectory clustering of the tumor-bearing dataset.** Related to **Fig.S4**, but trajectory clusters for the tumor-bearing condition. The "di" and "dfi" groups were further split into two. Manual clustering gives the seven tumor-bearing clusters shown in **Fig.3B**.

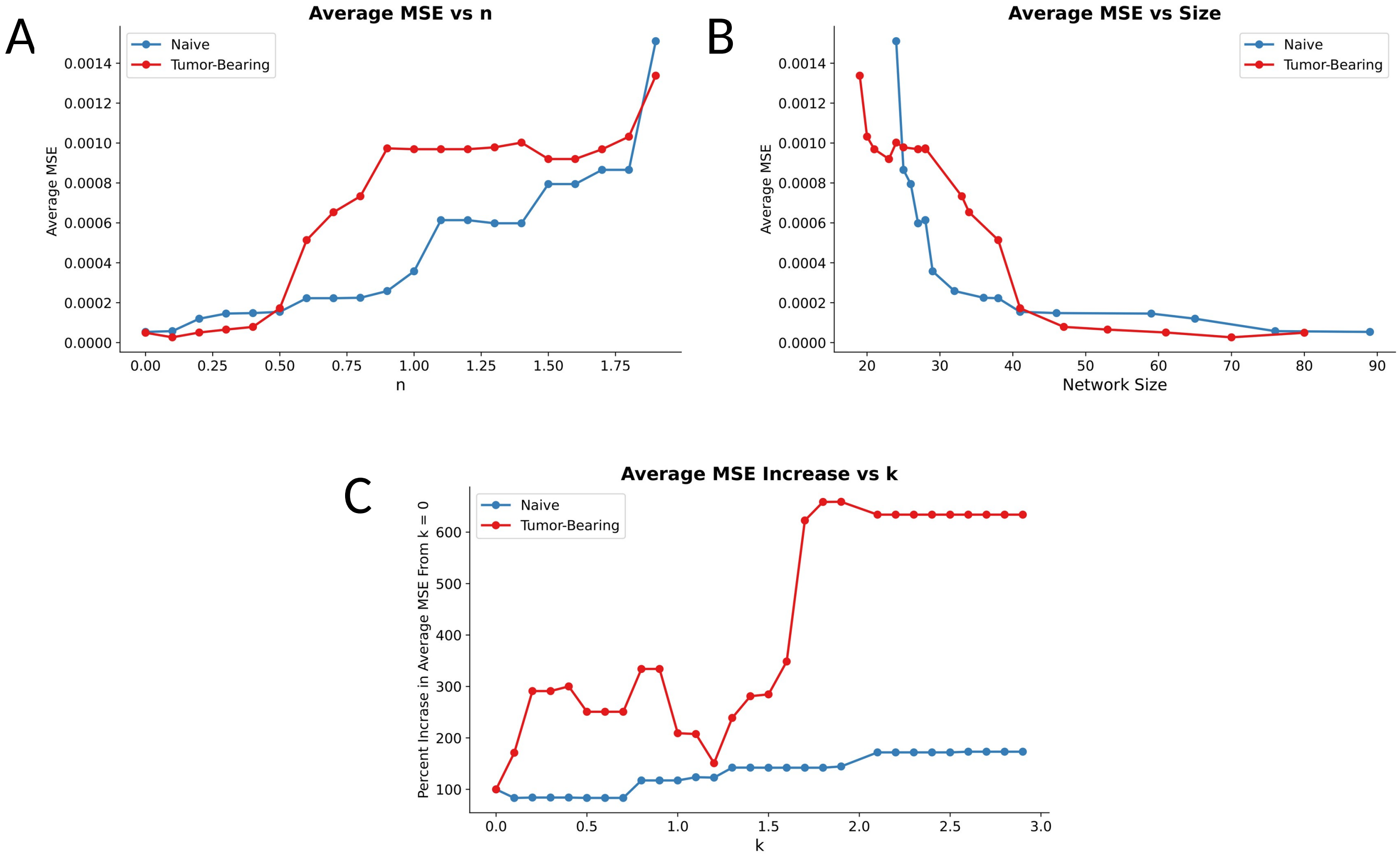


**Fig S6. Hyperparameter selection for neutrophil networks.** **(A)** Average MSE of the final optimized ODE models against the deletion parameter n, holding k = 1. **(B)** The same errors, plotted against optimized network size; error rises steeply beyond n = 0.5, corresponding to roughly 40 edges. **(C)** Percent increase in average MSE relative to k = 0 against the overlap penalty k, holding n = 1.

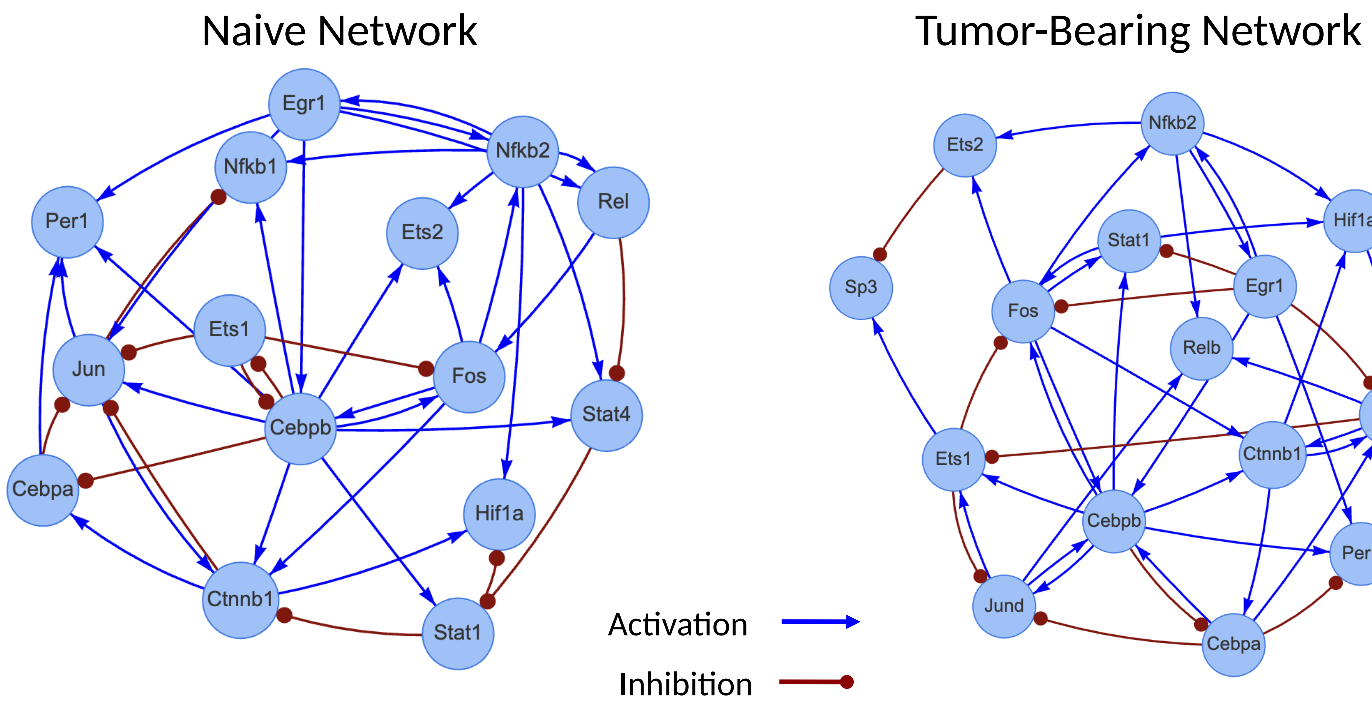


**Fig S7. Optimized naive and tumor-bearing networks shown separately.** The complete naive and tumor-bearing optimized TF regulatory networks. Blue lines with arrowheads represent activation, and red lines ending in dots represent inhibition. The edges associated with shared TFs appear overlaid with color-coded edges in **Fig.4A**.

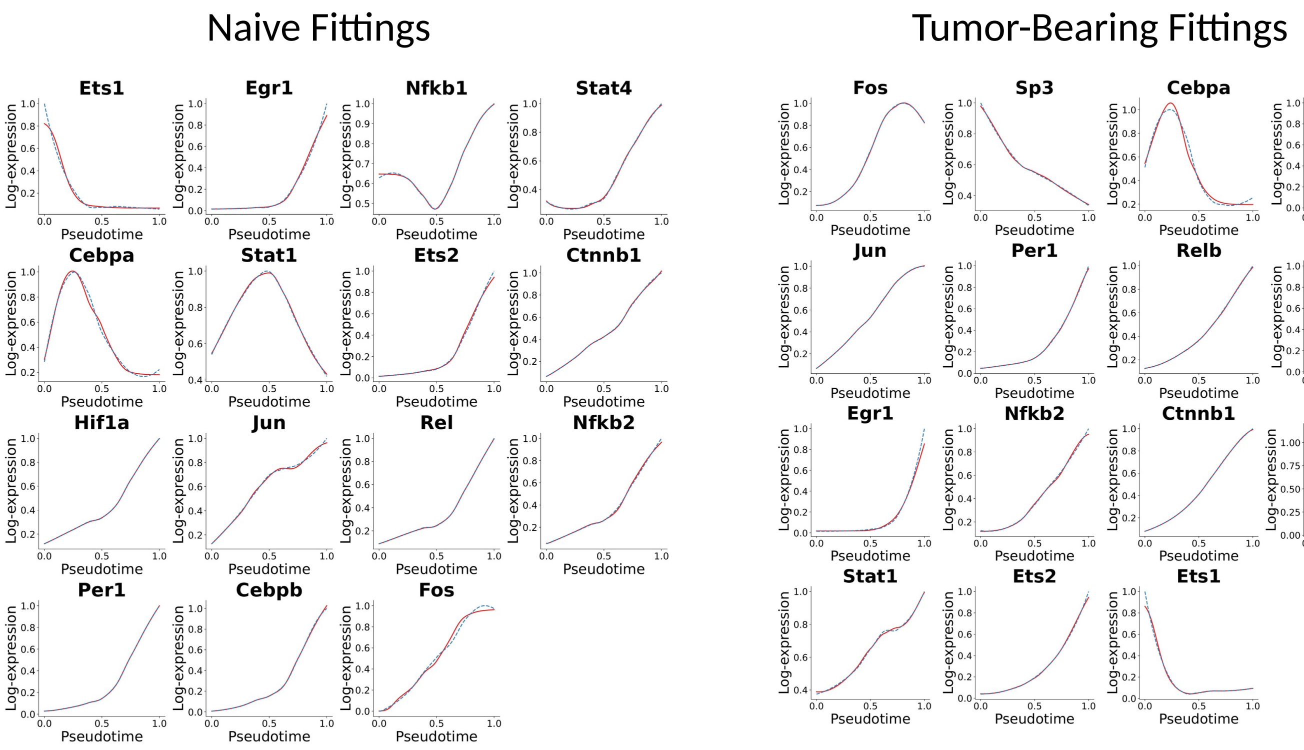


**Fig S8. Complete ODE fitting results for all network TFs.** Fitted trajectories for every TF in the naive and tumor-bearing optimized networks. Blue dotted lines are the smoothed scRNA-seq data and red solid lines are the optimized ODE model. The models track the data closely across all TFs in both conditions; selected panels appear in **Fig.4C**.

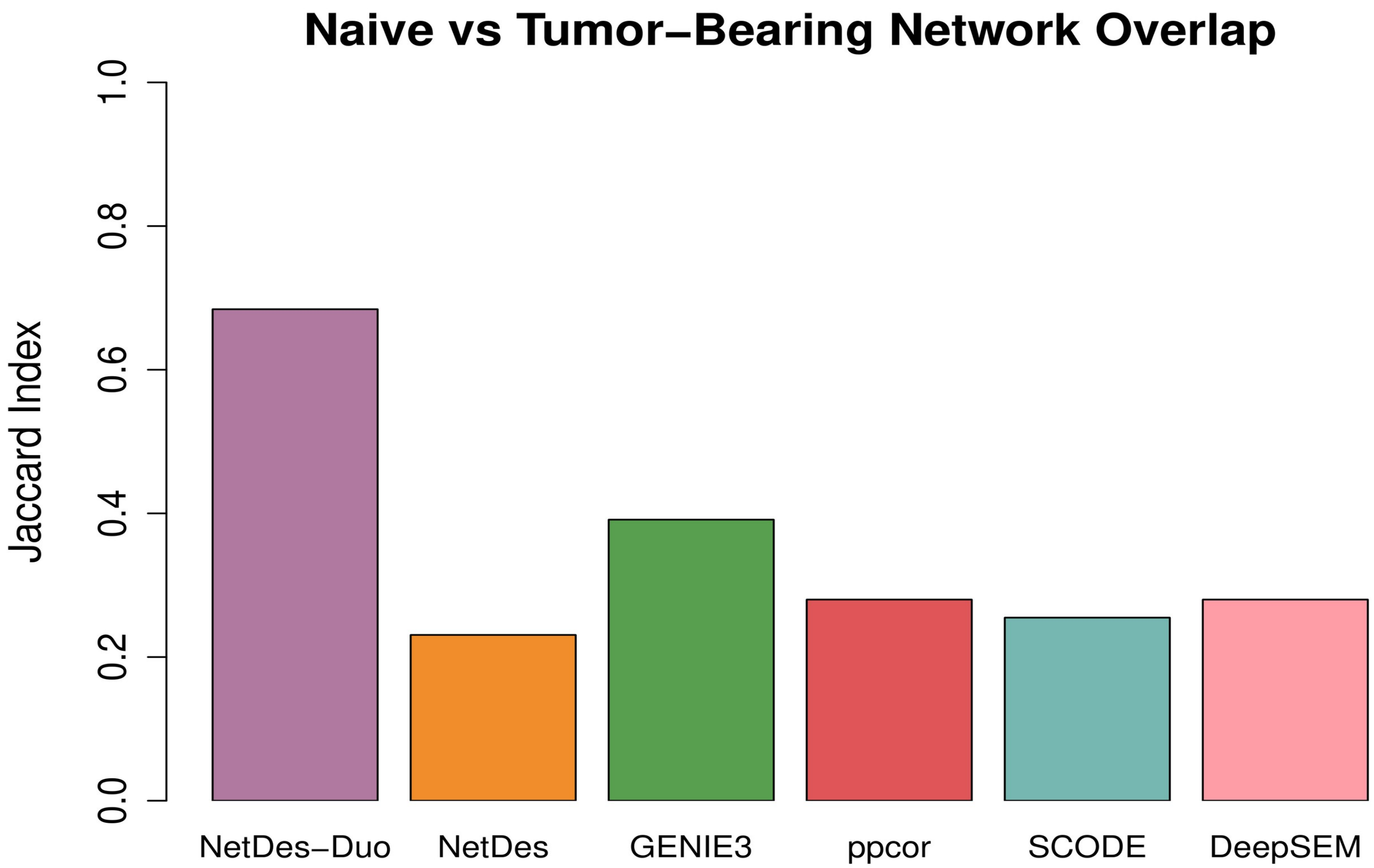


**Fig S9. Agreement between neutrophil networks by method.** Jaccard index between the inferred naive and tumor-bearing networks, for NetDes-Duo and the five comparison methods. Each comparison method was applied to the two conditions separately, with its network size matched to that of the corresponding NetDes-Duo network.

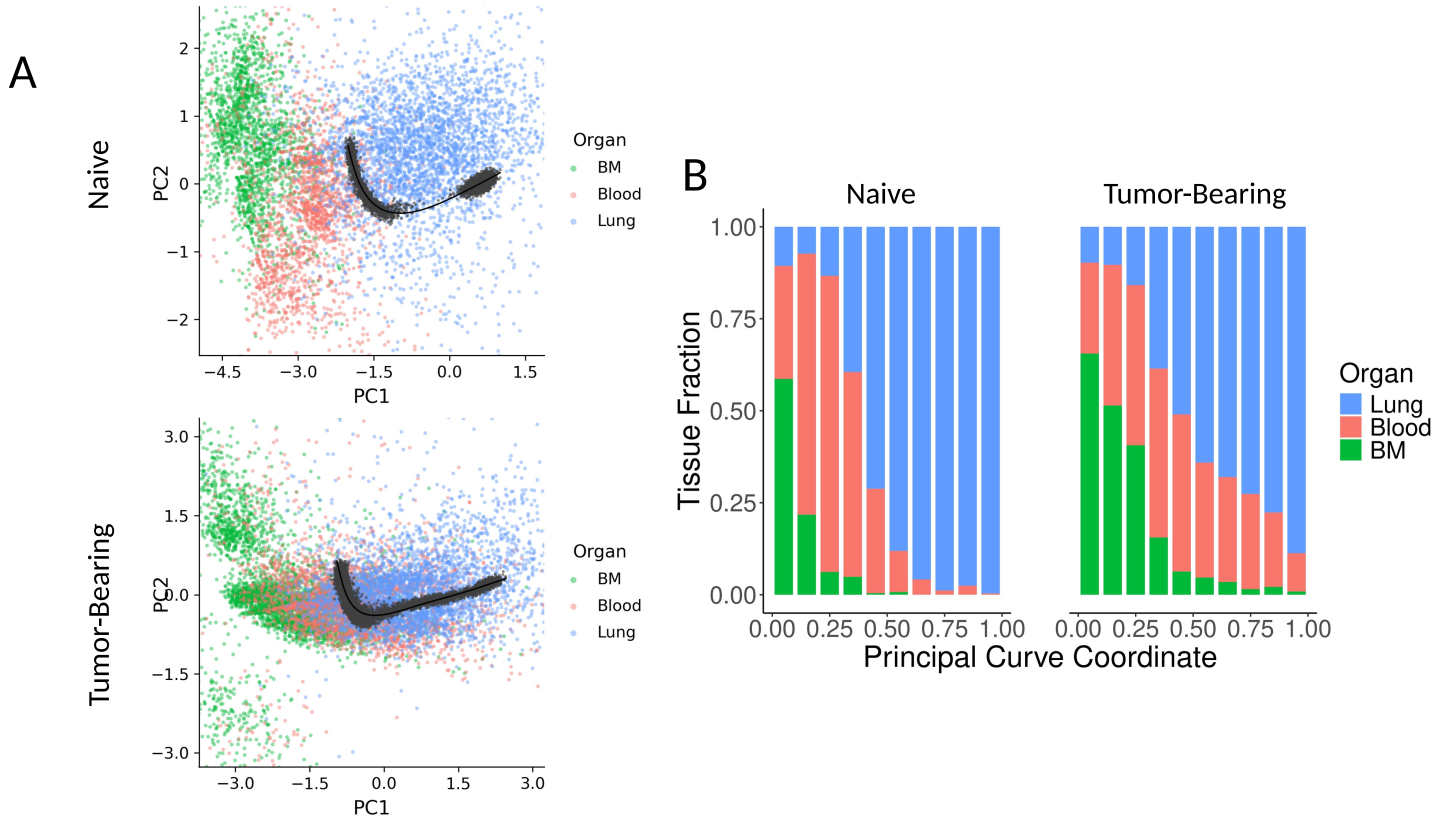


**Fig S10. Projection of scRNA-seq data onto the principal components of simulated data. (A)** Experimental gene expression profiles projected onto the first two principal components of simulated data, with the fitted principal curve in black, naive (top) and tumor-bearing (bottom). **(B)** Cells are binned by their principal curve coordinates. The stacked bar plots (naive on the left, tumor-bearing on the right) show the percent of measured cells in each bin contributed by each tissue with cells progressing from immature to mature as bins go from left to right.

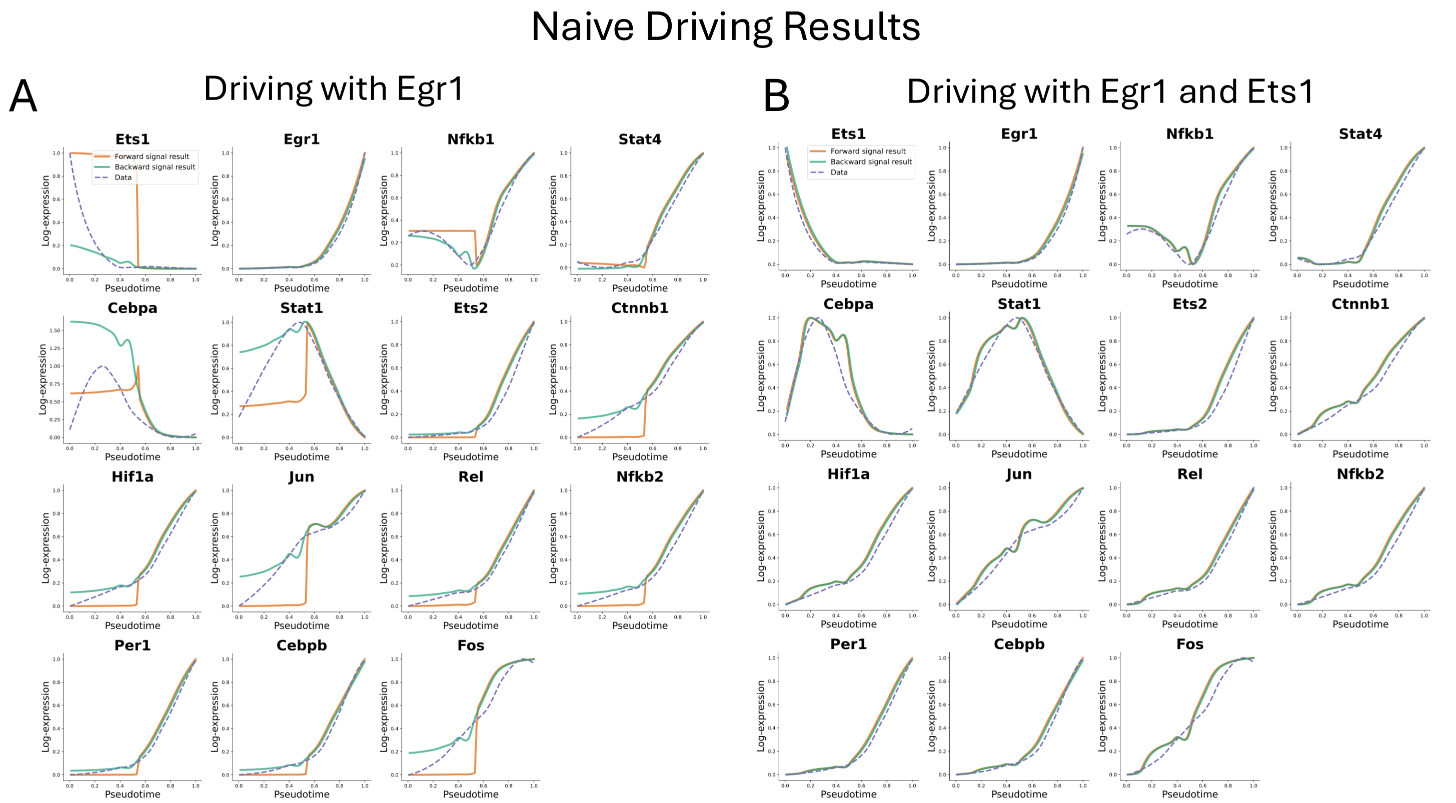


**Fig S11. Driving simulation results for the naive network.** **(A)** All TF responses when the naive network is driven by *Egr1* alone. **(B)** All TF responses when the naive network is driven by both *Egr1* and *Ets1*; selected curves are shown in **Fig.6C**. Orange curves show the forward driving result, green curves the backward driving result, and blue dashed curves the smoothed scRNA-seq data.

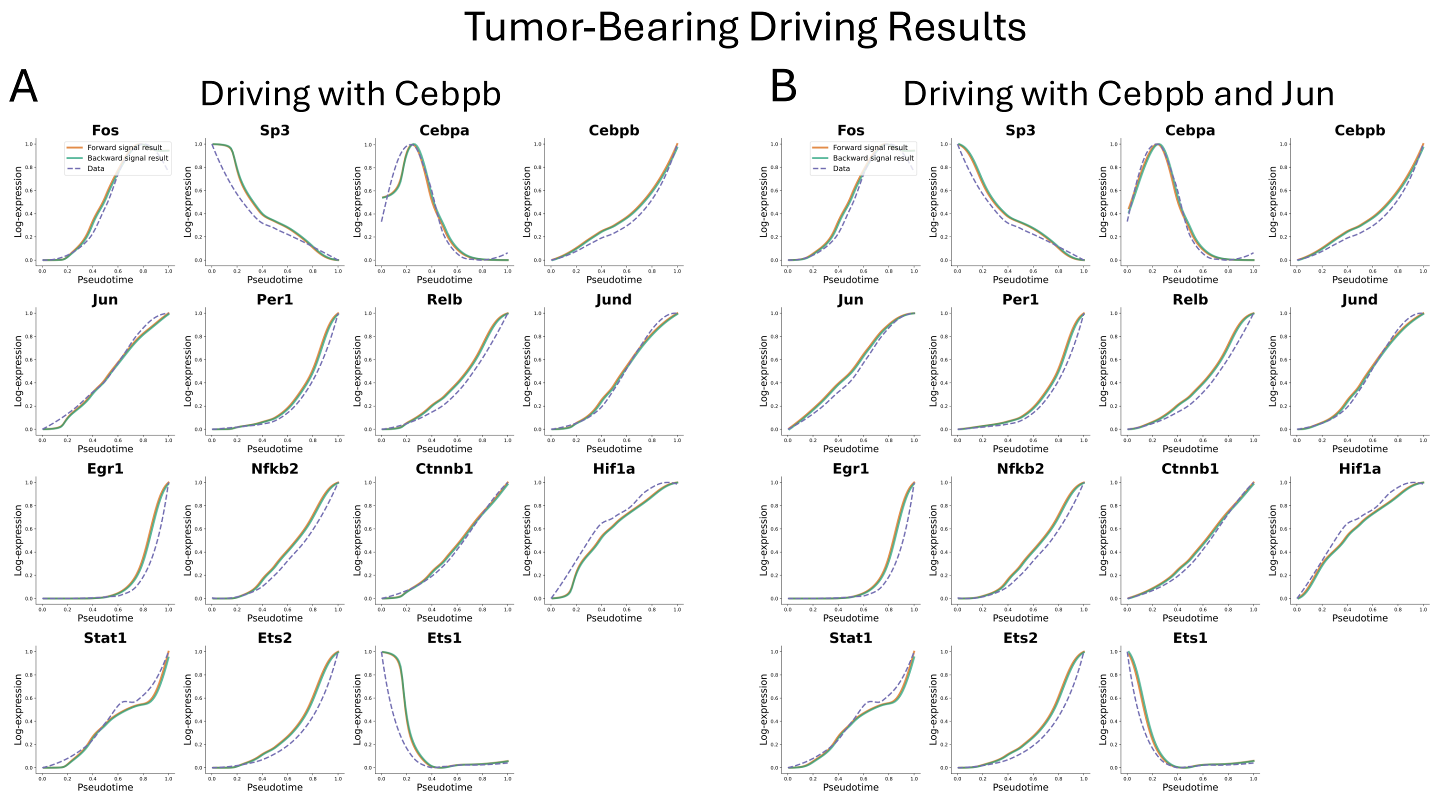


**Fig S12. Driving simulation results for the tumor-bearing network.** **(A)** All TF responses when the tumor-bearing network is driven by *Cebpb* alone. **(B)** All TF responses when the tumor-bearing network is driven by both *Cebpb* and *Jun*; selected curves are shown in **Fig.6C**.
